# Test-retest reliability of resting-state fMRI functional connectivity: impact of scan length and number of participants

**DOI:** 10.64898/2026.03.31.715533

**Authors:** Beatriz Vale, Marta M. Correia, Patrícia Figueiredo

## Abstract

Resting-state functional MRI has been widely used to study brain connectivity, yet the test-retest reliability of commonly used metrics remains a concern. To improve reliability, extended scan lengths and larger subject cohorts are often recommended. However, these solutions can be impractical and pose challenges, particularly in studies of clinical populations. Here, we systematically assess the reliability of two main types of functional connectivity measures: node-based connectome metrics (edge-level intraclass correlation coefficient [ICC], connectome-level ICC, fingerprinting, and discriminability); and voxel-based resting-state networks (RSNs) (spatial similarity of independent component analysis [ICA] derived RSN maps quantified using the Dice coefficient). Using data from the Human Connectome Project, we evaluated the effects of scan length (3.6, 7.2, 10.8, and 14.4 minutes) and number of participants (10, 20, 50, and 100), on both within-session and between-session reliability. We found that multivariate connectome metrics demonstrated greater reliability than edge-level measures, and that scan length had stronger influence on test-retest reliability than the number of participants. For connectome metrics, 14 minutes of scanning and a cohort of approximately 10 participants were sufficient to achieve reliable estimates. In contrast, RSN measures benefited from larger cohort sizes. Our findings provide practical guidelines for designing resting-state fMRI studies in terms of scan length and number of participants, balancing reliability and feasibility. Ultimately, protocol choices should be guided by the specific study objectives and the functional connectivity metric of interest.

**KEY POINTS:**

I. This work systematically evaluates the within- and between-session test-retest reliability of node-based and voxel-based resting-state fMRI functional connectivity metrics as a function of scan length and number of participants.
II. Longer scan length (∼11-14 minutes) improved the reliability of connectivity metrics, with specific requirements depending on the metric.
III. The number of participants had minimal effect on node-based metrics, but voxel-based metrics benefited from larger samples.

## 1. INTRODUCTION

The consistency of functional magnetic resonance imaging (fMRI) measurements upon repetition, known as test-retest reliability, is essential for the interpretation and validity of neuroimaging studies and directly impacts their statistical power and hence the ability to detect effects of interest [1], [2]. However, quantifying the reliability of resting-state fMRI measures of the brain’s functional connectivity (FC) [3] is complex, especially given the wide range of analysis methods and FC measures that can be employed, which can be roughly divided into node-based connectome [4], [5] and voxel-based resting-state networks (RSNs) [6], [7] approaches. Significant efforts have been made to develop appropriate reliability metrics and optimise them through choices in the preprocessing pipeline and study design [8], [9], [10]. Reliability metrics range from univariate measures (e.g., focused on individual functional connections or edges [11]) to multivariate approaches (e.g., the stability of whole-brain connectivity patterns, or fingerprinting [12], [13] or the spatial similarity between network maps [14], [15]).

For node-based connectome approaches, the intraclass correlation coefficient (ICC) [16] is commonly used to quantify the test-retest reliability of individual functional connections, or edges [15], [17], [18]. However, many studies have consistently reported poor test-retest reliability at the edge level [11], [19]. In contrast, multivariate reliability metrics, such as functional connectome fingerprinting [13] (based on correlations between connectomes), and discriminability [9] (based on distances between connectomes), have generally demonstrated higher reliability [20], [21]. These multivariate approaches suggest that differences in brain connectivity may be more reliably detected at the whole-connectome level rather than at the level of single connections [20].

For the identification of voxel-based RSNs [22], [23], a well-established approach consists of decomposing the fMRI data into statistically independent components using independent component analysis (ICA) and subsequently identifying RSNs based on the spatial similarity between their spatial maps and templates of so-called canonical RSNs [23]. Prior research indicates that ICA-derived RSNs show moderate test-retest reliability, although reliability values differ considerably across networks [10], [14], [24].

Scan length is a key factor influencing the reliability of resting-state FC estimates [8], [25], [26]. While studies have consistently shown that longer scans generally improve reliability [12], [18], [26], recommendations for sufficient scan length vary widely, ranging from as little as 5 minutes [27], [28] to over 90 minutes [29], [30], [31]. These discrepancies may stem from differences in the type of FC measures analysed and the respective reliability metrics used. Importantly, longer runs pose practical challenges for participants, including discomfort, fatigue, and increased head motion, all of which can reduce overall data quality.

Furthermore, the number of participants in the study is also crucial for determining the reliability of FC estimates [32], [33]. Although relatively few studies have directly examined its impact [34], [35], the broader fMRI literature consistently highlights the benefits of large cohorts for improving statistical power and reproducibility in the detection of group effects [36]. Statistically, larger cohorts reduce error variance, therefore increasing the likelihood of detecting true effects and reducing the incidence of false positives. Despite these benefits, practical constraints related to recruitment, cost, and time consumption [37] have limited the number of participants in neuroimaging research.

Thus, a trade-off is necessary. While both longer scan lengths and larger cohorts can contribute to improved reliability, overemphasising either can make studies impractical and compromise feasibility. Other factors impacting reliability include MRI acquisition parameters and preprocessing strategies [38], [39], [40], [41]. For example, FC reliability has been reported to improve when subjects have their eyes open during scanning [25], [42], whereas some preprocessing choices, such as global signal regression, can reduce reliability [8], [20].

Despite extensive research into the test-retest reliability of resting-state fMRI, the fundamental questions of how long to scan and how many participants to include to achieve sufficiently reliable resting-state FC estimates remain unclear. Here, we aim to fill this gap by systematically assessing the effects of scan length and number of participants on various test-retest reliability metrics of both connectome and RSN-based approaches. For this purpose, we use test-retest data from the Human Connectome Project (HCP) and assess test-retest reliability both within and between sessions. Specifically, we will assess the test-retest reliability of node-based functional connectomes using edge-level ICC and multivariate measures (connectome-level ICC, fingerprinting and discriminability) and the test-retest reliability of voxel-based RSNs using the spatial similarity between ICA-derived RSN maps [23]. Ultimately, we aim to provide recommendations on sufficient scan length and number of participants to optimise test-retest reliability of resting-state fMRI FC measures.

## 2. MATERIALS & METHODS

### 2.1. SUBJECTS AND DATA ACQUISITION

We used data from the Human Connectome Project (HCP), WU-Minn Consortium (http://www.humanconnectome.org/) [43], [44]. The sample consisted of 100 healthy, unrelated subjects (54 females), with most participants aged between 20 and 35 years (99 subjects) and one participant aged over 35 years. Each subject underwent two resting-state fMRI acquisitions on separate days, labelled as rfMRI_REST1 and rfMRI_REST2 in the HCP dataset. For each scanning session, data were acquired in two runs, using opposite phase encoding directions: left-to-right (LR) and right-to-left (RL).

Participants were instructed to lie still with their eyes open, fixate on a white cross displayed on a dark background, think of nothing in particular, and avoid falling asleep. Data were collected using a 3T Siemens Connectome Skyra MRI scanner with a 32-channel head coil. Functional images were acquired using a multiband gradient-echo EPI imaging sequence with the following parameters: 2 mm isotropic voxels, TR = 720 ms, TE = 33.1 ms, flip angle = 52°, field of view = 208 × 180 mm^2^, and a multiband factor of 8. A total of 1200 volumes were acquired, corresponding to a total scan length of 14 minutes and 24 seconds. For further acquisition details, refer to Smith et al. [45]. All participants provided written informed consent in accordance with the HCP protocols.

### 2.2. DATA PREPROCESSING

We used the minimally preprocessed HCP dataset [45], [46]. The preprocessing pipeline included artefact removal, motion correction, EPI distortion correction, alignment of individual structural scans, and registration to MNI152 standard space. Additionally, images were intensity normalised to a global mean, bias field correction was applied, and non-brain voxels were masked out. Full details on the preprocessing pipeline can be found in Glasser et al. [46].

In addition to the HCP minimal preprocessing, we carried out two additional preprocessing pipelines using the FMRIB Software Library (FSL) [47]. (i) *Minimal Pipeline*: high-pass temporal filtering with a cut-off frequency of 0.01 Hz to remove slow drift fluctuations, and spatial smoothing using a Gaussian kernel with a full width at half maximum (FWHM) of 3 mm. As a general rule of thumb, the FWHM was set to 1.5 times the voxel size [48]; and (ii) *Full Pipeline*: all steps in the minimal pipeline plus nuisance regression of motion parameters using the Friston 24-parameter model (6 head motion parameters, their first temporal derivatives, and the 12 corresponding squared items, totalling 24 motion regressors), as well as mean white matter (WM) and cerebrospinal fluid (CSF) BOLD time series (obtained by averaging the BOLD time series within WM and CSF masks derived from the FreeSurfer segmentation provided with the HCP dataset). The main results reported in the manuscript were obtained with the *Full Pipeline*; the comparison between the full and minimal pipelines is presented in the supplementary material.

### 2.3. TEST-RETEST RELIABILITY ANALYSIS

We assessed both within-session reliability, i.e. between the two runs within the same session [15], [49], and between-session reliability, i.e. between the two sessions [21], [50]. For the within-session reliability, we considered the two runs of session 1, i.e., REST1_LR and REST1_RL runs. For the between-session reliability, to avoid confounds between test-retest effects and phase encoding effects, we combined runs with the two different phase encodings evenly distributed across subjects: for half of the subjects, we compared REST1_LR with REST2_RL, and for the other half REST1_RL with REST2_LR.

A schematic overview of the methodological pipeline is presented in Figure 1.

**Figure 1.**
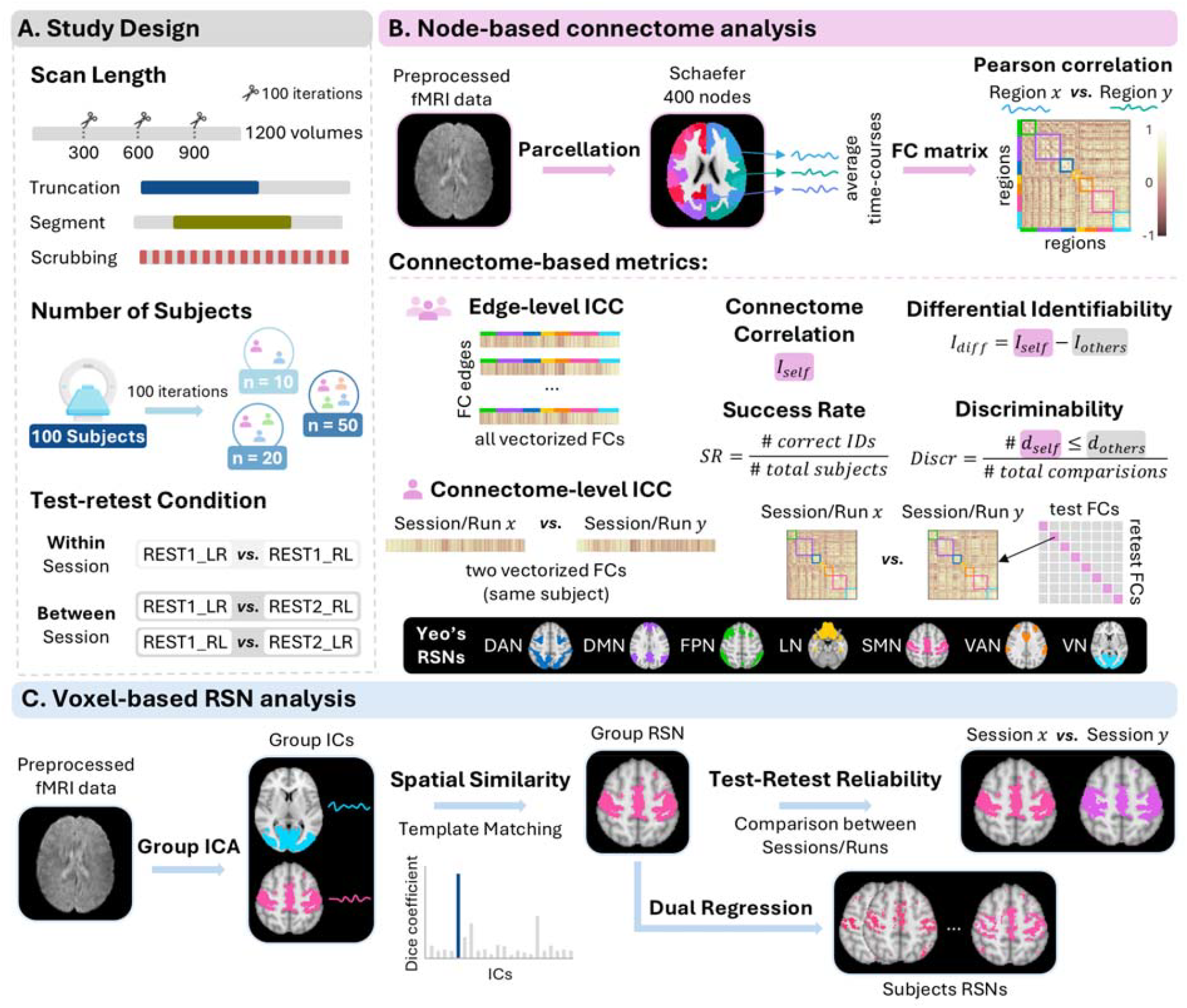
Schematic overview of the analysis pipeline. **(A) Study Design**. Two parameters were systematically varied: scan length and number of participants. For scan length, four numbers of volumes were tested (300, 600, 900 and 1200) using three strategies: (i) truncation, retaining only the initial portion of the scan; (ii) segment, extracting a continuous segment starting at a random time point and iii) scrubbing, removing a randomly selected subset of time points and concatenating the remainder data (100 iterations each). The number of participants was randomly subsampled from the full cohort of 100 to subsets of 10, 20 and 50 subjects (100 iterations each). For within-session comparisons, data from REST1_LR and REST1_RL were used; for between-session comparisons, data from runs of different sessions (REST1_LR vs REST2_RL and REST1_RL vs REST2_LR) were analysed. **(B) Node-based connectome analysis**. Preprocessed resting-state fMRI data were parcellated into 400 cortical regions using the Schaefer atlas, which corresponds to seven intrinsic networks: dorsal attention (DAN), default mode (DMN), fronto-parietal (FPN), limbic (LN), somatomotor (SMN), ventral attention (VAN) and visual (VN). FC matrices were computed by calculating the Pearson correlation between the average time courses of each pair of regions. The lower triangular portion of each FC matrix (excluding the diagonal) was then vectorised to calculate edge-level intra-class correlation (ICC) within each of the seven RSNs and the connectome-level ICC. The multivariate metrics (connectome fingerprinting and discriminability) were derived from comparisons of all FC matrices across subjects and sessions/runs. Fingerprinting was implemented using an identifiability matrix, computed as the Pearson correlation between all pairs of vectorised FC matrices. Discriminability was calculated using a Euclidean distance matrix across all subjects and sessions. **(C) Voxel-based RSN analysis**. Pre-processed resting-state fMRI data underwent group-level ICA. ICA-derived networks were compared to Yeo’s seven canonical RSNs, and the independent component (IC) with the highest Dice coefficient for each RSN was selected. For test-retest reliability, ICA-derived network maps were compared across sessions/runs using the Dice coefficient. Dual regression was performed to obtain subject-specific RSN spatial maps, which allowed for the same analysis to be performed based on subject-specific maps.

#### 2.3.1. NODE-BASED CONNECTOME ANALYSIS

Node-based FC analysis was performed using MATLAB (The MathWorks Inc., MA, USA version R2025a) (Figure 1B). To obtain functional connectomes, we parcellated the preprocessed fMRI data using the Schaefer atlas [5] with 400 parcels, by averaging the voxelwise BOLD time series within each parcel. This parcellation directly corresponds to the seven intrinsic networks defined by Yeo et al [23]. For each parcel, the time series were demeaned and bandpass-filtered between 0.01 and 0.1 Hz using a second-order Butterworth filter.

Symmetrical FC matrices were generated by computing the Pearson correlation coefficients between all pairs of regions. Pairwise FC values were then normalised using Fisher’s r-to-z transformation. For each subject and session, the lower triangular portion of the FC matrix (excluding the diagonal) was vectorised to represent the individual connectome as a single vector.

#### 2.3.2. Edge-level metrics

We assessed the reliability of FC at the edge level using the intraclass correlation coefficient (ICC) [11], [16], [20], a widely used metric that quantifies the consistency of a given measurement relative to the variability observed across participants. Specifically, we used ICC (2,1) to estimate the absolute agreement in single measurements, modelling sources of error as random (repeated sessions or runs). This model decomposes the total variance in FC values for a given edge into between-subject variance 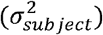, between-session/run variance 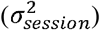, and residual error variance 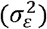 such that:

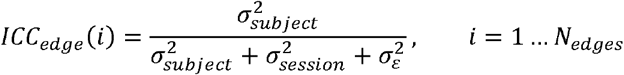

This formula quantifies the proportion of total variance attributable to true between-subject differences, with values ranging from 0 to 1. ICC values are commonly interpreted as: poor (< 0.4), fair (0.4 to 0.59), good (0.6 to 0.74), and excellent (≥ 0.75) [51]. These values were then grouped according to the seven intrinsic RSNs to which each edge belongs, and the average ICC across all edges of each RSN was reported.

#### 2.3.3. Connectome-level metrics

For the connectome level, we considered several different metrics that have been previously used in the literature.

##### Connectome-level ICC

To assess the reliability of each subject’s overall FC profile across sessions/runs, we computed ICC for the vectorised FC matrices [52], [53]. Here, ICC(2,1) is applied per subject. ICC is computed across the vectorised FC matrix from each session/run of the same subject, treating edges as the unit of observation. The model decomposes the total variance in FC values into between-edge variance 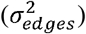, between-session/run variance for the same individual 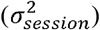, and residual error variance 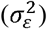, such that:

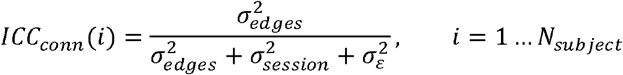

This quantifies how consistent an individual’s whole-connectome FC pattern is across sessions. One ICC value was computed per subject, and the average across all subjects was reported.

##### Connectome correlation

For each subject, we calculated the Pearson correlation between the vectorised FC matrices of the two runs or sessions. The within-participant connectivity matrix correlation reported was then obtained by averaging these correlations across all subjects.

##### Fingerprinting metrics

The fingerprinting approach evaluates whether individual connectomes are more similar within a subject across sessions than between different subjects [13], [54]. We evaluated this based on two different metrics: differential identifiability and fingerprinting success rate.

We implemented the framework introduced by Amico & Goñi [12], who proposed the concept of differential identifiability (*I*_*diff*_). An identifiability matrix *A* was constructed, where the elements represent the Pearson correlation between the FC matrices of test and retest sessions across all subjects. *I*_*diff*_ of the population is defined as the difference between self-identifiability (*I*_*self*_), calculated as the average Pearson correlation between the two sessions of the same subject (i.e., the average of the main diagonal elements of *A*), and *I*_*others*_, defined as the average Pearson correlation between sessions of different subjects (i.e., the average of the off-diagonal elements of the matrix *A*):

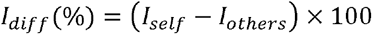

A higher *I*_*diff*_ indicates a greater ability to identify individuals based on their FC patterns.

We also computed the fingerprinting “success rate” as defined by Finn et al. [13]. Considering the identifiability matrix *A*, a subject was considered correctly identified if their *I*_*self*_ (diagonal elements of *A*) exceeded the correlation with every other subject’s retest session (same row) and every other subject’s test session (same column). The success rate of an individual was computed as the proportion of such comparisons in which *I*_*self*_ was the highest. The overall success rate is the mean of these individual rates across all subjects.

##### Discriminability

Discriminability (*Discr*) is a nonparametric multivariate reliability metric introduced by Bridgeford et al. [9], which quantifies the extent to which within-subject similarity exceeds between-subject similarity based on distances between FC matrices (rather than their Pearson correlation).

We generated a pairwise Euclidean distance matrix *B* between all scans (i.e., all sessions across all subjects), where each element *B*(*i,j*) represents the Euclidean distance between scan *i* and scan *j*. For each scan, its within-subject distance (*d*_*within*_, distance to the other session of the same subject) was compared against its distance to every session of every other subject individually (*d*_*between*_). Discriminability is defined as the proportion of these pairwise comparisons in which the within-subject distance was smaller than or equal to the between-subject distance, summed across all scans and normalised by the total number of comparisons made as follows:

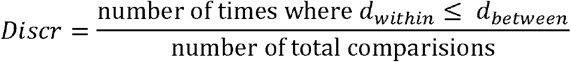

#### 2.3.4. VOXEL-BASED RSN ANALYSIS

For the voxel-based RSN analysis (Figure 1C), group-level ICA was performed separately for each run of each session (REST1_LR, REST1_RL, REST2_LR and REST2_RL) using FSL’s MELODIC [47]. The number of components was fixed at 50, representing an intermediate value that aims to balance the risks of overfitting or underfitting data [55]. The z-statistic spatial maps of each independent component (IC) were binarised using a threshold of z > 3. Furthermore, dual regression was performed to obtain subject-specific IC time series and spatial maps.

Firstly, the spatial similarity between each IC spatial map and the templates of the seven canonical RSNs proposed by Yeo et al. [23] was computed using the Dice coefficient, to identify the ICA-derived RSNs as the 7 IC’s yielding the maximum Dice coefficient with the respective RSN template [23][56]. Specifically, each IC was classified as one of the following RSNs: dorsal attention network (DAN), default mode network (DMN), frontoparietal network (FPN), limbic network (LN), somatomotor network (SMN), ventral attention network (VAN), or visual network (VN). Dual regression was then performed using FSL’s command *dual_regression* to obtain the map of each RSN at the subject level.

An exception was made for the VN due to its well-documented subdivision into multiple subcomponents in ICA (e.g., medial, lateral and occipital visual areas following Smith et al. [22]), typically yielding 3 ICs with similarly high Dice coefficients with the VN template. In this case, the IC selected was not always the one with the highest Dice coefficient. Instead, the three ICs with the top Dice coefficients were examined by visual inspection, and the ones corresponding to the same visual subnetwork across sessions/runs were chosen (as illustrated in **Figure 2**Figure 2). Specifically, the ICs with the highest Dice coefficients from the three datasets under consideration (e.g., REST1_LR, REST1_RL and REST2_RL) were examined. If all three corresponded to the same VN subnetwork, the IC with the highest Dice coefficient was kept. However, if the top-ranked IC in one dataset represented a different VN subnetwork (e.g., as in **Figure 2**Figure 2, IC1 of REST1_LR), the second-ranked IC was selected instead, provided it also showed a high Dice coefficient and a closer anatomical match to the other two sessions. Overall, this adjustment was necessary in 16% of cases.

**Figure 2.**
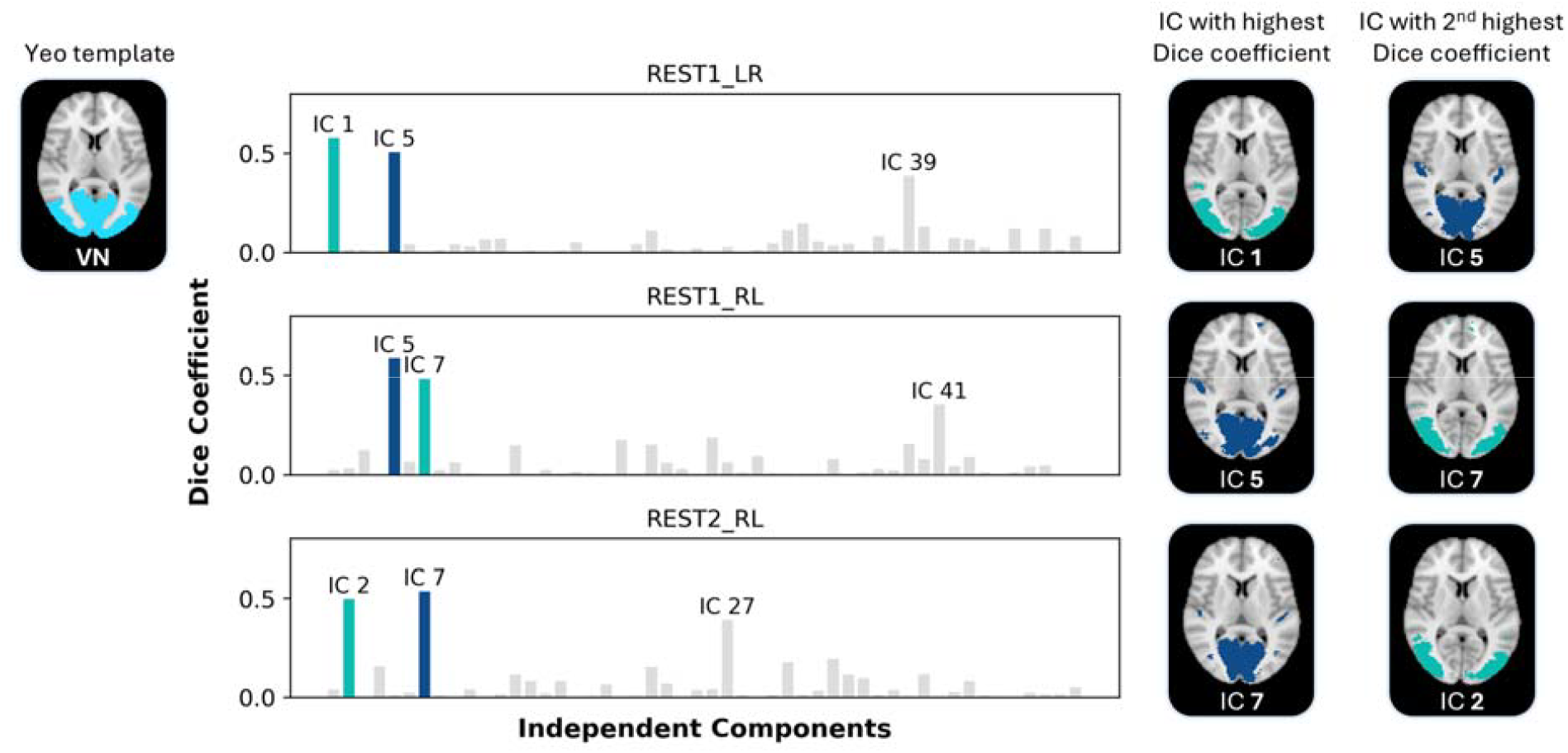
Selection of the visual RSN independent component (IC): illustrative example. For each dataset (REST1_LR, REST1_RL, and REST2_RL), bar plots show the Dice coefficients between each IC map and the canonical visual network (VN) template from Yeo et al. [23] (shown on the top left, in cyan). The values of the 3 highest Dice coefficients are shown on top of the respective bars. The chosen IC is represented in dark blue, the second possible choice IC in green, and all other ICs in grey bars. The spatial maps of the ICs with the highest and 2^nd^ highest Dice coefficients are shown on the right. In this example, for the REST1_LR session, the IC with the highest Dice coefficient (IC 1) was not chosen. Instead, the IC with the second-highest Dice coefficient (IC 5) was selected because it provided a more consistent anatomical match to the VN identified in the other two datasets. This approach ensures valid spatial test-retest reliability analysis across all runs, accounting for the VN’s known subdivision into multiple components.

Finally, the test-retest reliability of the ICA-derived RSN spatial maps was assessed by computing the Dice coefficient between runs (within-session) or sessions (between-session). This analysis was performed both on group RSN maps and the subject-specific maps.

#### 2.3.5. INFLUENCE OF SCAN LENGTH AND NUMBER OF PARTICIPANTS ON RELIABILITY

To evaluate the influence of scan duration on the test-retest reliability metrics, we systematically varied the number of volumes included in the analysis (Figure 1A). For this purpose, we considered a temporal truncation approach whereby scans were truncated to include only the initial portion of the time series, corresponding to: 300 volumes (3 minutes 36 seconds), 600 volumes (7 minutes 12 seconds), 900 volumes (10 minutes 48 seconds) and the full 1200 volumes (14 minutes 24 seconds).

For the connectome-based analysis, we further tested two other strategies to manipulate scan length besides temporal truncation (*truncation*), yielding the same number of volumes (300, 600, 900 and 1200). The second strategy consisted of random segment extraction (*segment*), whereby contiguous segments of data were extracted starting from randomly chosen time points. The third strategy (*scrubbing*) consisted of removing a randomly selected subset of time points from the full time series and subsequently concatenating the remaining volumes. For both the random segment extraction and scrubbing approaches, each analysis was repeated 100 times with a random selection of the initial time point or the subset of excluded time points, respectively. To assess the influence of the number of participants (Figure 1A), we randomly selected a subset of the total number of subjects: 10, 20, 50, and the complete 100 subjects. For cohort sizes smaller than the full cohort of 100 subjects, 100 iterations were performed, with subjects randomly selected in each iteration. For the ICA-based measures, only 10 iterations were performed for each combination of scan length and number of participants due to computational demands.

#### 2.3.6. STATISTICAL ANALYSIS

All statistical analyses were conducted in R (version 4.5.2; https://www.r-project.org/). For each reliability metric, a linear model was fitted with scan length (300, 600, 900, and 1200), number of participants (10, 20, 50, 100), and retest condition (within- vs. between-session) as fixed factors, including a scan length × number of participants interaction. For metrics derived at the RSN level (ICC edge-level, voxel-based metrics), RSN was additionally included as a fixed factor in all interactions. Main effects and interactions were assessed using ANOVA. To evaluate the effect of the temporal downsampling strategy (*truncation, segment, scrubbing*) on reliability, a separate set of models was fitted using a fixed sample of 100 participants, with scan length, strategy, and their interaction as fixed factors. Similarly, to compare preprocessing pipelines, the pipeline was included alongside scan length and number of participants as fixed factors.

Full statistics are provided in Supplementary Tables S1-S6. Post hoc pairwise comparisons were conducted using Bonferroni correction to adjust for multiple comparisons. “Sufficient” scan length and cohort size were defined as the lowest value beyond which no further significant pairwise differences were observed.

## 3. RESULTS

All results reported here are based on the *Full Preprocessing* pipeline. A detailed comparison with the *Minimal Preprocessing* pipeline can be found in the supplementary material.

### 3.1. NODE-BASED CONNECTOME ANALYSIS

For node-based connectome metrics, the test-reliability was assessed as a function of scan length, number of participants, retest condition and temporal downsampling strategy. For clarity, we presented the results in two separate figures, emphasising different factors. Figure 3 shows the effects of scan length and number of participants for the truncation downsampling strategy. Figure 4 shows the effects of scan length and temporal downsampling strategy on the node-based connectome reliability metrics, using a fixed sample of 100 subjects, four scan lengths (3.6 to 14.4 minutes) and three temporal downsampling methods (*truncation, segment, scrubbing*).

**Figure 3.**
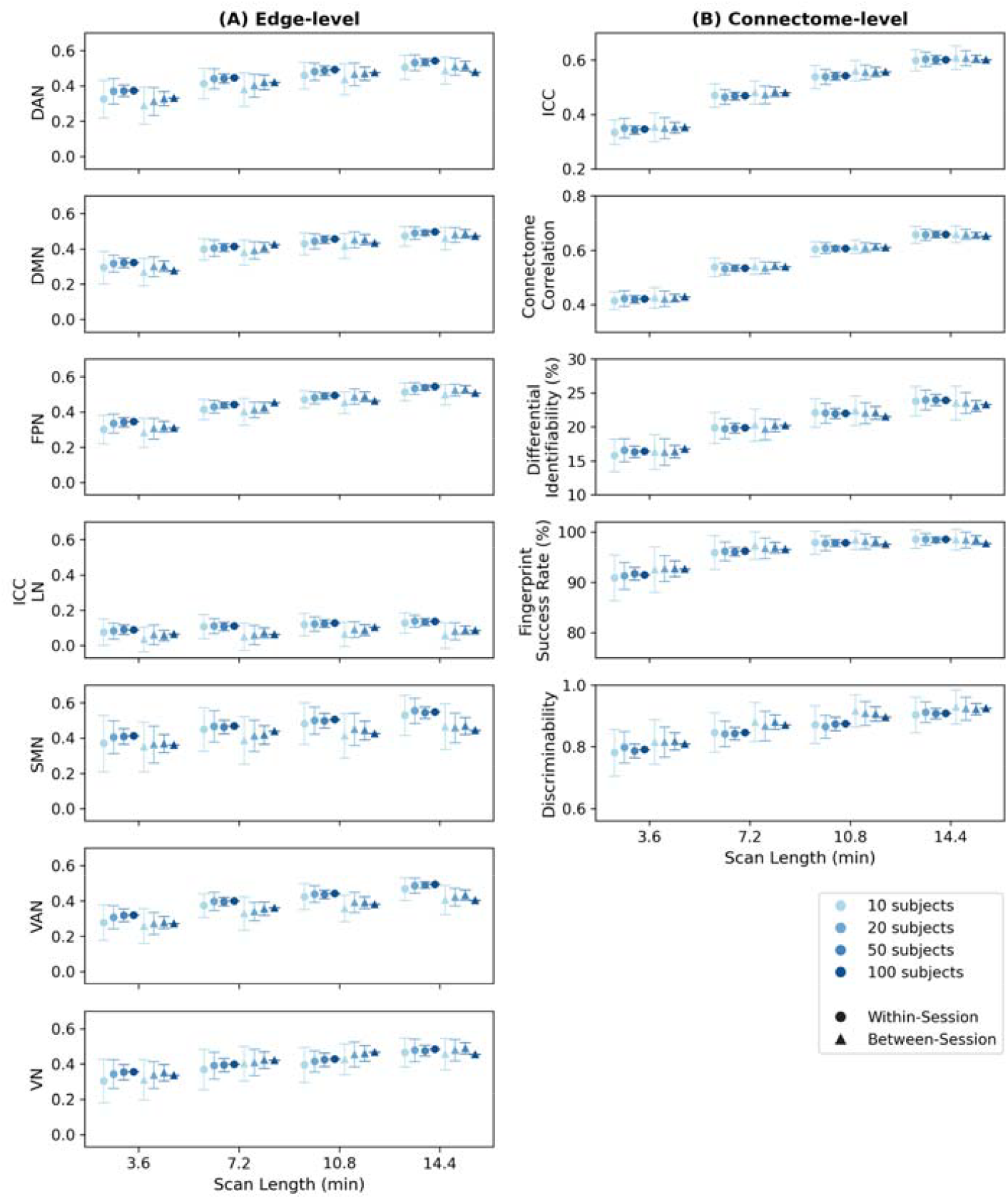
Test-retest reliability of node-based connectomes, for edge-level (A) and connectome-level (B) metrics, as a function of scan length (in minutes) and number of participants, for the full preprocessing pipeline. (A) Edge-level ICC per RSN. (B) Connectome-level ICC, connectome correlation (), differential identifiability (), fingerprinting success rate and discriminability. Dots represent the mean and error bars the standard deviation of the mean across 100 iterations (except for the case of 100 subjects, for which only 1 iteration is performed). Scan lengths correspond to 300 (3 minutes 36 seconds), 600 (7 minutes 12 seconds), 900 (10 minutes 48 seconds) and 1200 (14 minutes 24 seconds) volumes, respectively. Data points are slightly offset along the x-axis for visual clarity. DAN: dorsal attention network, DMN: default mode network, FPN: fronto-parietal network, LN: limbic network, SMN: somatomotor network, VAN: ventral attention network, VN: visual network.

**Figure 4.**
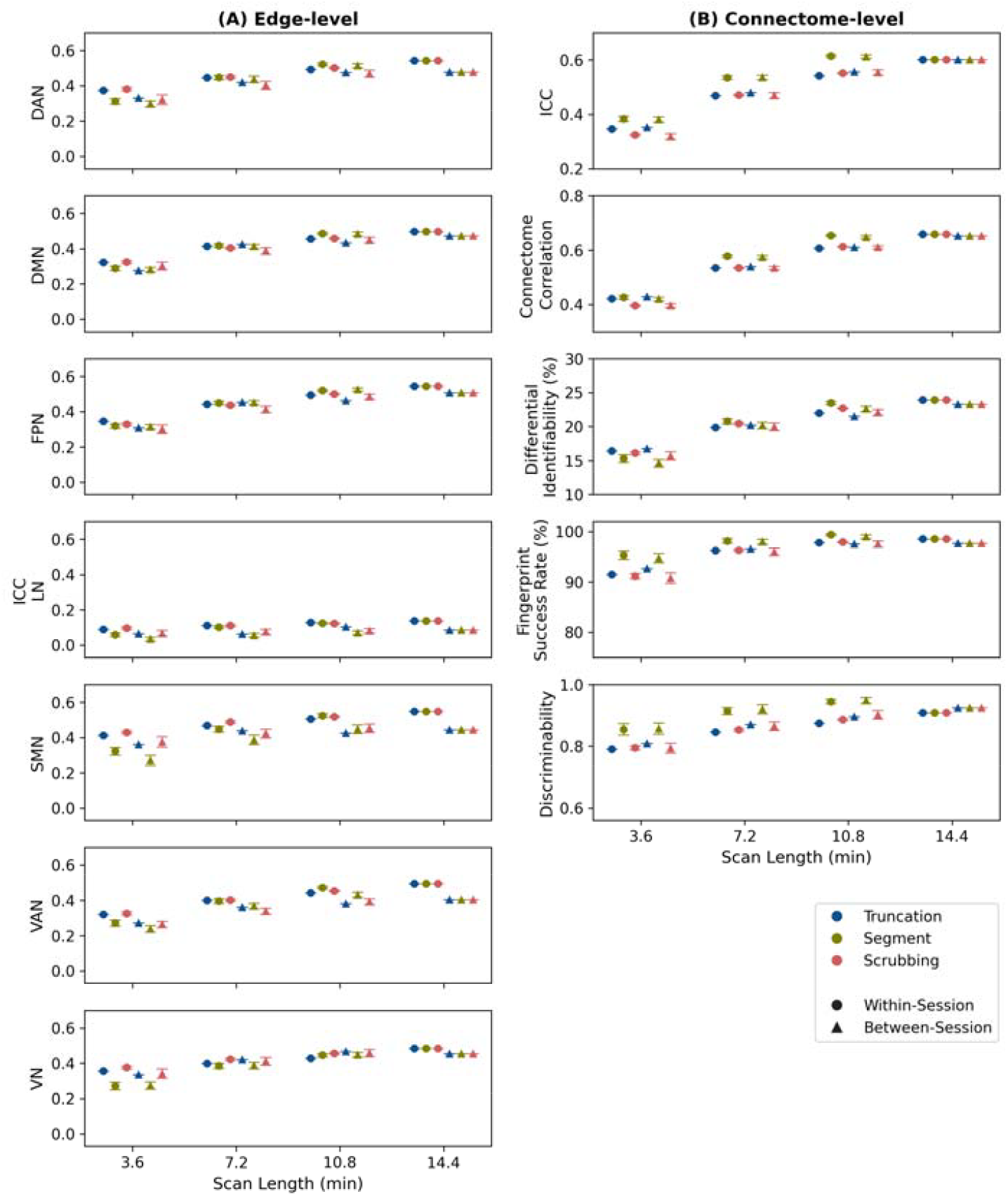
Effect of temporal downsampling strategy on the test-retest reliability of node-based connectome metrics, as a function of scan length (in minutes), for a fixed sample of 100 subjects, and the full preprocessing pipeline. (A) Edge-level ICC: average of within-RSN edges per RSN. (B) Connectome-level ICC, connectome correlation (), differential identifiability (), fingerprinting success rate and discriminability. Blue curves (truncation) represent the inclusion of timepoints from the beginning of the scan. Green curves (segment) represent continuous segments of data extracted from randomly selected start points. Pink curves (scrubbing) represent the removal of randomly selected timepoints from the time series. Dots represent the mean and error bars the standard deviation of the mean over 100 iterations (except for truncation, for which only 1 iteration is performed). Scan lengths correspond to 300 (3 minutes 36 seconds), 600 (7 minutes 12 seconds), 900 (10 minutes 48 seconds) and 1200 (14 minutes 24 seconds) volumes, respectively. Data points are slightly offset along the x-axis for visual clarity. DAN: dorsal attention network, DMN: default mode network, FPN: fronto-parietal network, LN: limbic network, SMN: somatomotor network, VAN: ventral attention network, VN: visual network.

Across all metrics, scan length (Figure 3) had a strong positive effect on reliability, and longer scans consistently improved edge-level ICC, connectome-level ICC, within-participant connectome correlations, differential identifiability, fingerprinting success rate, and discriminability (Table S1A). However, the scan length at which further increasing the number of volumes no longer produced statistically significant results differed by metric. For fingerprinting success rate, reliability increased with longer scans and plateaued at 900 volumes (10.8 minutes), as no significant differences were observed between 900 and 1200 volumes (10.8 and 14 minutes, respectively). However, edge-level ICC, connectome-level ICC, connectome correlation, differential identifiability and discriminability all required the full scan length of 1200 volumes (14.4 minutes) to reach their highest reliability (Table S1B).

At the edge-level ICC (Figure 4A), ICC values increased with scan length but varied significantly across RSNs. To achieve fair reliability (0.40 < ICC < 0.59), the DAN, DMN, FPN, SMN, and VN required at least 900 volumes (10.8 minutes) of data, the VAN required the full 1200 volumes (14.4 minutes), and the LN did not reach fair reliability even at the full scan length. None of the RSNs reached good reliability (ICC ≥ 0.60) at the edge level.

In contrast, multivariate and whole-connectome metrics (Figure 4B) showed higher reliability overall. From 600 volumes (7.2 minutes) onward, connectome-level ICC values were in the fair range (ICC > 0.40), and good reliability (ICC ≥ 0.60) was reached only at the full scan length of 1200 volumes (14.4 minutes). At the full scan length, connectome-level metrics showed the highest values: connectome correlations ranged from ∼0.6 to 0.7, differential identifiability reached ∼23-25%, fingerprinting success rate exceed 97%, and discriminability surpassed 0.9.

The temporal downsampling method also had a statistically significant effect on all metrics except edge-level ICC (Figure 4, Table S5A). In general, random segment extraction produced the highest reliability, truncation the lowest, and scrubbing showed intermediate performance. Post hoc comparisons (Table S5B), however, revealed that differences between methods were not always significant for every metric: for connectome correlation, all temporal downsampling strategies differed significantly; for discriminability and fingerprinting success rate, there were no significant differences between scrubbing and truncation; and for differential identifiability, no significant pairwise differences between methods were observed. A significant interaction between scan length and downsampling method was observed for all metrics except edge-level ICC (Table S5C).

Since truncation is the most realistic approach for simulating different scan lengths, this method was selected to examine the combined effects of scan length and number of participants (Figure 3). As reported above, scan length had a statistically significant effect on all evaluated metrics (). In contrast, the number of participants did not show a significant main effect for any of the metrics evaluated (edge-level ICC, connectome-level ICC, correlation between within-participant connectivity matrices, differential identifiability, fingerprinting success rate and discriminability, Table S2A). No significant interaction between the number of participants and scan length for any of the metrics was observed (Table S2F). Figure 5 illustrates two examples of identifiability matrices across different scan lengths and numbers of participants, where it is possible to visually observe the impact of variability in the chosen subjects.

**Figure 5.**
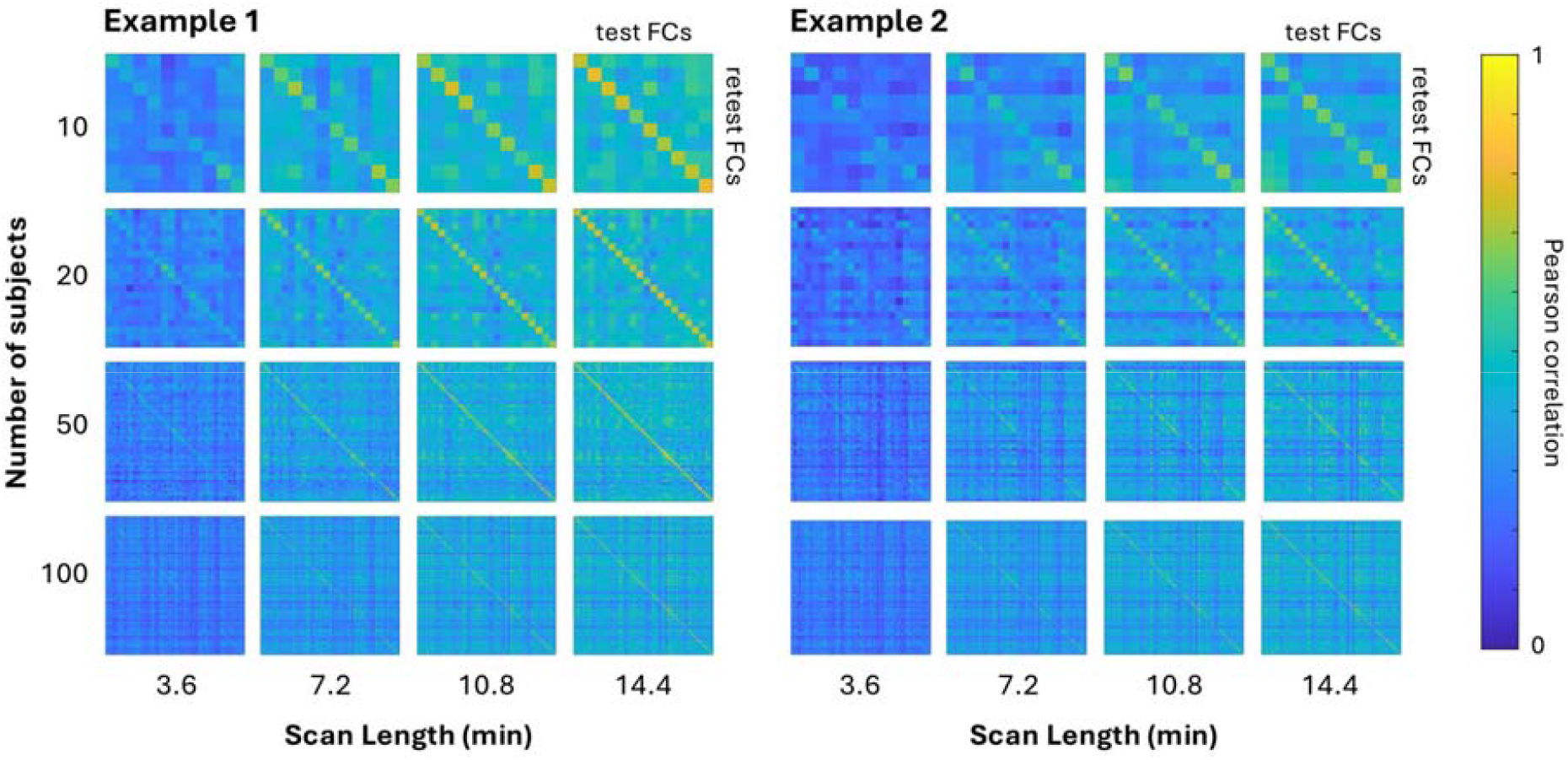
Two examples of identifiability matrices for fingerprinting test-retest analysis of node-based functional connectomes. Each matrix shows the Pearson correlation between test and retest functional connectivity (FC) matrices for one iteration of the between-session analysis. The subject subsets in example 1 and example 2 are distinct, but their relative ordering is consistent within each example matrix. Rows correspond to different numbers of subjects (10, 20, 50, 100 subjects) and columns correspond to different scan lengths: 300 (3 minutes 36 seconds), 600 (7 minutes 12 seconds), 900 (10 minutes 48 seconds) and 1200 (14 minutes 24 seconds) volumes, respectively.

We assessed the effect of the retest condition (within versus between sessions) (Table S3). For edge-level ICC, within-session comparisons showed significantly higher reliability than between-session comparisons. In contrast, connectome-level ICC and discriminability showed the opposite pattern, with significantly higher values for between-session acquisitions compared to within-session acquisitions.

Finally, the preprocessing pipeline had a significant effect on all connectome-based metrics except fingerprint success rate (Figure S3, Table S6). Metrics consistently showed higher values when processed using the full pipeline compared to the minimal pipeline, suggesting that additional preprocessing steps, such as nuisance regression of motion parameters and WM and CSF mean time series, improved reliability across all metrics.

### 3.2. VOXEL-BASED RSN ANALYSIS

At the group level maps, Dice coefficients were relatively stable across scan lengths. For spatial test-retest reliability, which assesses the similarity of ICA-derived RSNs across sessions/runs, scan length had no significant main effect, indicating that as little as 300 volumes (3.6 minutes) of data were sufficient (Figure 6A, Table S1A). Similar results were observed for spatial similarity, which quantifies the overlap between ICA-derived networks and Yeo’s canonical RSNs (Figure S1, Table S1A).

**Figure 6.**
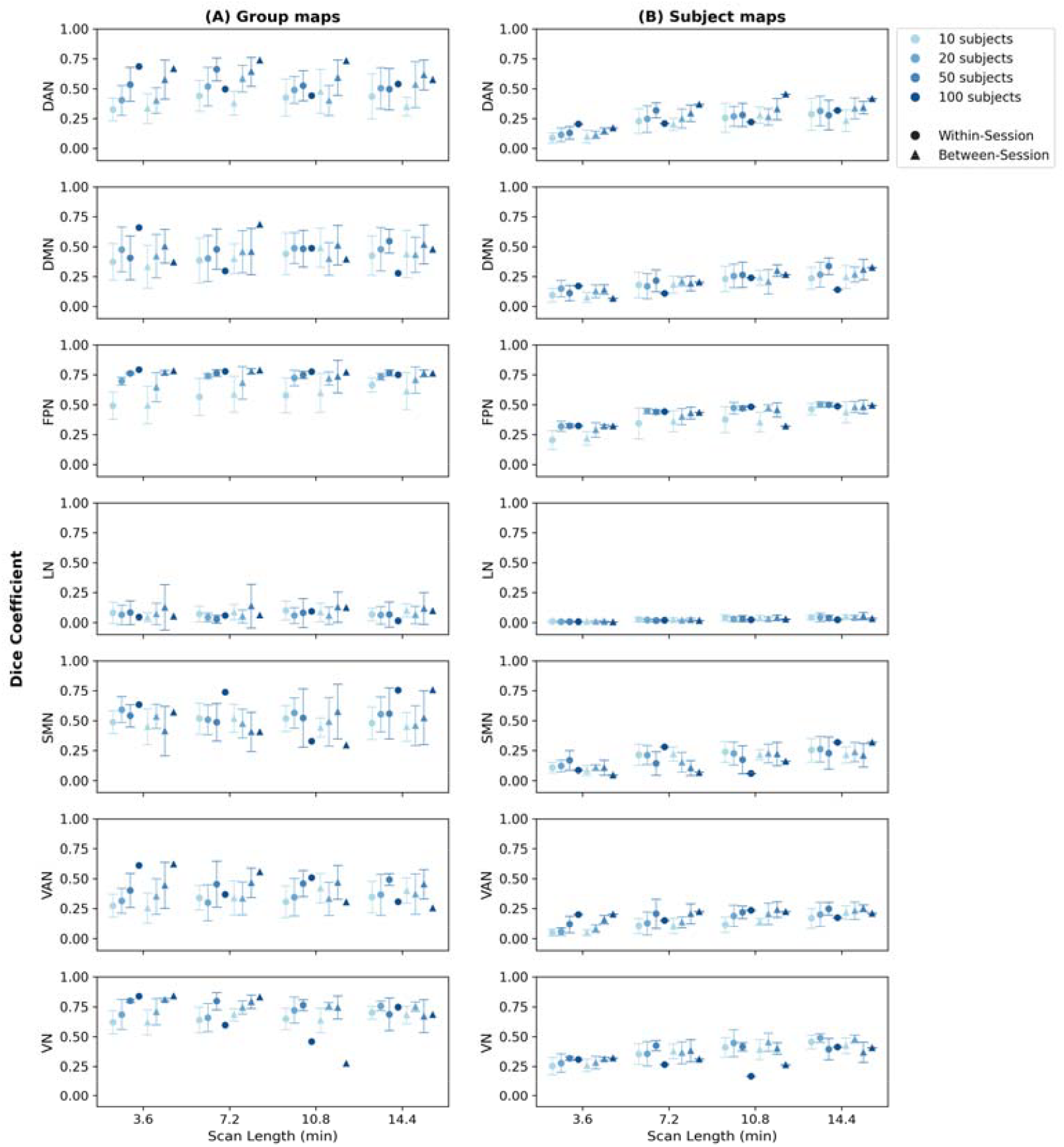
Test-retest reliability of voxel-based RSNs, for group maps (A) and subject maps (B), as a function of scan length (in minutes) and number of participants, for full preprocessing pipeline. All plots represent Dice coefficients for each RSN, between runs (within-sessions) and between sessions. Dots represent the mean and error bars the standard deviation across 10 iterations (except for the case of 100 subjects, for which only 1 iteration is performed). Scan lengths correspond to 300 (3 minutes 36 seconds), 600 (7 minutes 12 seconds), 900 (10 minutes 48 seconds) and 1200 (14 minutes 24 seconds) volumes, respectively. Data points are slightly offset along the x-axis for visual clarity. DAN: dorsal attention network, DMN: default mode network, FPN: fronto-parietal network, LN: limbic network, SMN: somatomotor network, VAN: ventral attention network, VN: visual network.

At the subject level maps, the effect of scan length was more pronounced. For both spatial test-retest reliability (Figure 6B, Table S1B) and spatial similarity (Figure S2, Table S1B), all post hoc pairwise comparisons between scan lengths were significant, and the full 1200 volumes (14.4 minutes) scan yielded the highest Dice coefficients.

The number of participants had a significant main effect on Dice coefficients at only the group level maps but not when considering the subject-specific maps (Table S2A). At the group level, spatial reliability (Figure 6A) generally increased with the number of participants, and differences between cohort sizes were statistically significant (Table S2A). The same patterns were observed for the spatial similarity with the RSN templates (Figure S1). For spatial reliability, post hoc comparisons indicated that Dice coefficients were significantly different across all cohort sizes tested until 50 subjects (Table S2B). Although mean Dice values tended to increase with more participants, some specific scan lengths and RSNs (e.g., group maps of the DMN, VAN at 14.4 minutes within-session) showed slightly lower Dice coefficients at 100 subjects compared with 50. For spatial reliability, no significant interaction between the number of participants and RSN was observed (Table S2C). Similarly, scan length and number of participants showed no significant interaction across all voxel-based RSN metrics, at either the group or subject level (Table S2F).

There were no statistically significant differences between within-session and between-session acquisitions, at either the group or subject level (Table S3).

Dice coefficients varied significantly across RSNs at both the group and subject levels (Table S4), with group-level values being consistently higher than subject-level ones for all RSNs. For spatial test-retest reliability, the FPN and VN showed the highest Dice coefficients. For spatial similarity, the DAN, DMN, SMN and VN showed the highest values. Overall, spatial test-retest reliability tended to yield higher Dice coefficients than spatial similarity.

A positive correlation was observed between RSN template size and mean Dice coefficient across all analyses. The LN, which had the smallest template, consistently exhibited the lowest Dice values across both metrics and analysis levels. In contrast, larger networks such as VN and DMN tended to show higher spatial overlap (Figure S5). This pattern suggests that Dice coefficients may be inherently sensitive to template size.

The preprocessing pipeline had no statistically significant main effect on spatial reliability (Figure S4, Table S6).

## 4. DISCUSSION

In this study, we investigated the test-retest reliability of node-based connectome metrics (i.e., edge- and connectome-level ICC, differential identifiability, and discriminability) and ICA-derived networks under varying conditions, including scan length and number of participants, as well as test-retest condition and temporal downsampling strategy. Our findings revealed four key observations. First, longer scan length improved the reliability of connectivity metrics, with 11-14 minutes of data being generally sufficient to achieve stable estimates, though the precise requirement varied depending on the metric examined. Second, the number of participants had limited influence on node-based connectome metrics, whereas ICA-derived metrics benefited from larger samples. Third, the retest condition varied depending on whether data was acquired within the same session or between sessions, with different metrics showing opposite patterns. Finally, while individual connections (edge-level) showed poor reliability, multivariate metrics demonstrated moderate to high reliability. ICA-derived networks exhibited moderate spatial test-retest reliability, although this pattern was not consistent across all RSNs.

### 4.1. EFFECT OF SCAN LENGTH

Univariate metrics such as edge-level ICC showed limited reliability (*ICC*_*edge*_ ≈ 0.4 - 0.5), depending on the RSN under study, largely aligning with prior HCP-based studies reporting mean ICC values ranging from ∼0.2 to ∼0.5 across full scan lengths [18], [57], [58], [59]. In our data, we observed that edge-level ICC improved with longer scan lengths but did not reach 0.6, the threshold for good reliability. Approximately 10 minutes of data were sufficient to achieve fair reliability (*ICC* > 0.4) for most RSNs, with the exception of the VAN and LN, and extending the scan length beyond this point did not yield meaningful improvements. However, for achieving high edge-level reliability, recent studies on precision resting-state fMRI approaches that combine high temporal resolution with extended acquisition durations (e.g., > 60 minutes) have reported ICC values in the range of 0.8 − 0.9 [30], [60]. In contrast, connectome-level ICC, which reflects the consistency of the entire connectivity matrices, showed higher reliability (*ICC*_*conn*_ ≈ 0.5), in line with previous studies [49], [57], [61]. As with edge-level ICC, connectome-level ICC increased with scan length, but in this case, at full-length, reached the threshold for good reliability.

Multivariate metrics, which take into account the full connectivity matrix, demonstrated the highest test-retest reliability as reported in previous studies [20], [62], [63], with values improving as the scan length increased. For differential identifiability, our results are partially consistent with Amico & Goñi (2018) [12], who reported that optimal identifiability increased as a function of the number of fMRI volumes, using the same dataset as in our study (HCP 100 unrelated subjects). Their PCA-based reconstruction approach yielded higher identifiability with 200 volumes than the value obtained at full length using the original FC matrices. This suggests that reconstruction techniques and preprocessing choices may be useful to improve identifiability even with shorter acquisitions. Similarly to *I*_*diff*_, the simpler measure of the correlation between within-participant FC matrices improved steadily with scan length, with the highest reliability achieved at full duration.

Regarding discriminability, to our knowledge, no prior study has directly investigated the effect of scan length. This is an important gap to address, as discriminability is a nonparametric multivariate metric that offers a complementary, distance-based approach to fingerprinting. Our results suggest that discriminability also benefits from longer acquisitions, reaching values > 0.8 at longer durations, which is within the range considered good by previous studies [21], [64].

Regarding ICA-derived RSNs, both the test-retest reliability and similarity to the templates significantly improved with increasing scan length, when assessing the RSN spatial maps at the subject level. In contrast, for group-level RSN maps, approximately 300 volumes (3.6 minutes) of data were sufficient to achieve stable and reliable estimates. This finding aligns with previous studies indicating that variations in pipelines can significantly affect the reliability of subject-level RSNs but have minimal influence on the reliability of RSNs at the group level [65]. Although longer scan lengths tended to increase the Dice coefficient and improve the spatial consistency of ICA-derived networks, the effect was not uniform across all RSNs. Prior studies have shown higher reliability in higher-order association networks such as DMN, DAN, VAN and FPN compared to sensory networks such as the VN and SMN, attributed to higher interindividual variability and lower within-subject fluctuations in cognitive networks [14], [66]. However, network reliability varies by analysis method as group-level reliability exceed subject-level reliability [65], as observed in our results. Additionally, previous studies have shown that sensorimotor networks (particularly somatomotor and auditory) showed better individual-subject reliability than some association networks [65]. Our results diverge from this sensory-network pattern in the sense that the VN was one of the most reliable networks in our analysis. However, we also observed a positive correlation between RSN template size and Dice coefficient. Larger networks such as VN, FPN, SMN and DMN showed higher spatial overlap, while the smallest network (LN) consistently exhibited the lowest values. This suggests that RSN reliability may depend on methodological factors such as template size, preprocessing pipeline, and IC dimensionality rather than functional network classification alone.

Overall, these findings reinforce that longer scan lengths tend to improve test-retest reliability across metrics, with the best reliability being reported at ∼11-14 minutes for most measures. Considering practical constraints such as participant fatigue, increased motion and reduced compliance associated with very long scans, a scan length of 11-14 minutes appears to provide a practical balance between data quality and feasibility for studies aiming to achieve reliable resting-state estimates.

### 4.2. EFFECT OF NUMBER OF PARTICIPANTS

For node-based metrics (edge-level ICC, connectome-level ICC, connectome correlation, differential identifiability, success rate and discriminability), even a number of participants of 10 yielded reliability estimates that were statistically comparable to those obtained with 100 participants. However, although the mean values for these smaller numbers of participants did not differ significantly from those of larger samples, the associated standard deviation across the 100 iterations was notably higher when fewer participants were included. This suggests that achieving reliable results with small samples may depend on the inclusion of particularly stable or “good” subjects, making such estimates sensitive to inter-individual variability as seen is Figure 5, where it is possible to observe how different sets of subjects yield different identifiability matrices.

This variability reflects the well-documented inter-individual differences in resting-state brain connectivity [23], [67] which can influence the stability of derived metrics. Larger numbers of participants, while not statistically improving mean reliability in our analysis, nonetheless provide a better representation of this variability and lead to more robust estimates across iterations [68]. In the case of univariate measures (such as edge-level ICC), which are, in nature, less reliable, a trend towards more participants was required for sufficient power to observe univariate effects [33]. However, the trade-off between statistical stability and practical feasibility should be considered, as increasing the number of participants poses logistical and financial demands associated with recruiting, scanning and retaining large cohorts. Given these constraints and the results of the present study, a number of participants of approximately 20 appears to provide a reasonable balance between data stability and resource requirements for studies employing similar connectivity-based metrics.

If the study primarily focuses on ICA-derived RSNs, our findings indicate that a number of participants of approximately 50 provides the most robust results, yielding higher spatial test-retest reliability across runs and with the RSNs templates. This suggests that, for ICA-based analyses, larger cohorts help stabilise network estimates and improve test-retest reliability. It is important to acknowledge, however, that this conclusion is based on analyses performed with a fixed number of independent components (50 ICs), which may have influenced the observed outcomes.

### 4.3. WITHIN VS. BETWEEN-SESSION RELIABILITY

Measurements performed within a session or between sessions comprise different sources of variance in fMRI data, naturally affecting the test-retest reliability of fMRI-derived metrics. In our study, we observed differential effects across metrics. Edge-level ICC showed significantly higher reliability for within-session compared to between-session data. These findings partly align with previous studies, which have often reported higher reliability within-session repeated measures compared to between-session measures [26], [69]. Within-session comparisons avoid the confound of participant repositioning and benefit from consistent scanner settings and experimental conditions. In contrast, both discriminability and fingerprint success rate exhibited the opposite pattern, with higher values for between-session acquisitions. This unexpected observation deserves further investigation.

### 4.4. EFFECT OF TEMPORAL DOWNSAMPLING METHOD

We investigated the effect of temporal downsampling methods to better understand whether reliability was primarily driven by longer acquisition times or by the number of available time points, focusing on node-based connectome metrics. Based on prior studies, such as Shah et al. [18], who used the HCP 500 subjects dataset and suggested that reliability depends more on the effective length of the scan than the number of time points used in the analysis, we anticipated that scrubbing (randomly removing time points) would have minimal impact on reliability, while truncation (retaining only the initial time points) would result in lower reliability. However, contrary to this expectation, our findings did not support this pattern. This divergence may reflect differences in parcellation choices or analysis approaches as they used seed-based connectivity from selected ROIs, whereas we used node-based FC analysis.

Interestingly, our results indicate that random segment extraction consistently outperformed both truncation and scrubbing. Since resting-state connectivity metrics ultimately reflect patterns of synchrony in brain activity, they may be particularly sensitive to the mental state of the participant during specific periods of the scan. This finding might suggest that continuous segments of data may better preserve the temporal dynamics necessary for stable connectivity estimation, even if the total acquisition time is shorter. One potential explanation is that truncation, which always takes the start of the scan, may capture a period when the brain has not yet reached a steady state for resting-state connectivity estimation, thereby reducing reliability.

From a practical perspective, neither scrubbing nor random segment extraction truly corresponds to a shorter scan length in a real-world acquisition, but these findings provide valuable insight into how data sampling affects test-retest reliability. An alternative approach for investigating how the number of data points could influence reliability may involve adjusting temporal resolution. For example, Cassone et al. [38] demonstrated how manipulating the repetition time (TR) can influence the reliability of functional fingerprinting.

### 4.5. LIMITATIONS

First, the generalisability of our findings is limited by the use of the Human Connectome Project dataset, which consists of high-quality data acquired from healthy young adults under standardised acquisition protocols. These results may not directly extend to clinical populations or studies performed under less controlled conditions. Future work should assess whether the same scan length and number of participants recommendation holds in more heterogeneous setups or cohorts.

Second, we accessed scan lengths up to 14 minutes only. While for several metrics this appeared sufficient, some measures continued to improve up to 14 minutes, suggesting that longer acquisitions could provide additional benefits. Additionally, a methodological consideration is that both within-session and between-session comparisons involved different phase encoding directions (LR and RL). While we balanced phase encoding directions across subjects for between-session analysis to minimise confounding, this introduces a source of protocol variation that may influence reliability estimates.

For ICA-derived metrics, a key limitation concerns the definition and identification of RSNs. As emphasised in a recent review [55], there is no universally accepted approach for RSN definition, which contributes to variability across studies. Moreover, the present analysis was performed with a fixed model order (50 independent components), which may have influenced the observed outcomes. Future research should evaluate how varying ICA model order influences the optimal number of participants and scan length to improve spatial similarity and test-retest reliability.

## 5. CONCLUSION

There has been a growing interest in the reliability of resting-state connectivity measures. In this study, we systematically examined the test-retest reliability of node-based connectome metrics and voxel-based RSNs under varying conditions of scan length and number of participants. Our results suggest that a scan length of about 11-14 minutes and a cohort of around 10 participants are sufficient for most connectivity-based metrics, whereas ICA-derived measures benefit from larger samples.

A key contribution of this work is the systematic evaluation of test-retest reliability of multiple commonly used node-based and voxel-based metrics across several experimental conditions simultaneously (scan length, number of participants, and retest condition), whereas previous studies have typically examined individual factors or metrics in isolation.

Ultimately, the choice of scanning protocol, both in terms of scan length and number of participants, should be guided by the specific goals of the study, with the aim of maximising reliability and validity while maintaining feasibility.

## Acknowledgements/Funding

This work was funded by FCT through LARSyS-FCT (DOI: 10.54499/LA/P/0083/2020). MMC is funded by UKRI MRC through unit grant SUAG/019 G116768.

## Conflicts of Interest

The authors declare no conflicts of interest.

## Data Availability Statement

The dataset used in this study is comprised of 100 unrelated subjects available in the Human Connectome Project (HCP) repository (http://www.humanconnectome.org/).

## Notes

http://www.humanconnectome.org/

### Competing Interest Statement

The authors have declared no competing interest.

### Summary of Updates

This version includes revisions throughout the manuscript and supplementary materials.

## REFERENCES

[1] M. Gell et al., “How measurement noise limits the accuracy of brain-behaviour predictions,” Nat. Commun., vol. 15, no. 1, p. 10678, 2024, doi: 10.1038/s41467-024-54022-6.

[2] X.-N. Zuo, T. Xu, and M. P. Milham, “Harnessing reliability for neuroscience research,” Nat. Hum. Behav., vol. 3, no. 8, pp. 768–771, 2019, doi: 10.1038/s41562-019-0655-x.

[3] B. Biswal, F. Z. Yetkin, V. M. Haughton, and J. S. Hyde, “Functional connectivity in the motor cortex of resting human brain using echo-planar MRI.,” Magn. Reson. Med., vol. 34, no. 4, pp. 537–541, Oct. 1995, doi: 10.1002/mrm.1910340409.

[4] J. D. Power et al., “Functional network organization of the human brain.,” Neuron, vol. 72, no. 4, pp. 665–678, Nov. 2011, doi: 10.1016/j.neuron.2011.09.006.

[5] A. Schaefer et al., “Local-Global Parcellation of the Human Cerebral Cortex from Intrinsic Functional Connectivity MRI,” Cereb. Cortex, vol. 28, no. 9, pp. 3095–3114, Sep. 2018, doi: 10.1093/cercor/bhx179.

[6] C. F. Beckmann, M. DeLuca, J. T. Devlin, and S. M. Smith, “Investigations into resting-state connectivity using independent component analysis.,” Philos. Trans. R. Soc. London. Ser. B, Biol. Sci., vol. 360, no. 1457, pp. 1001–1013, May 2005, doi: 10.1098/rstb.2005.1634.

[7] J. S. Damoiseaux et al., “Consistent resting-state networks across healthy subjects.,” Proc. Natl. Acad. Sci. U. S. A., vol. 103, no. 37, pp. 13848–13853, Sep. 2006, doi: 10.1073/pnas.0601417103.

[8] X.-N. Zuo et al., “Toward reliable characterization of functional homogeneity in the human brain: preprocessing, scan duration, imaging resolution and computational space.,” Neuroimage, vol. 65, pp. 374–386, Jan. 2013, doi: 10.1016/j.neuroimage.2012.10.017.

[9] E. W. Bridgeford et al., “Eliminating accidental deviations to minimize generalization error and maximize replicability: Applications in connectomics and genomics,” PLOS Comput. Biol., vol. 17, no. 9, p. e1009279, Sep. 2021, [Online]. Available: 10.1371/journal.pcbi.1009279

[10] W. R. Shirer, H. Jiang, C. M. Price, B. Ng, and M. D. Greicius, “Optimization of rs-fMRI Pre-processing for Enhanced Signal-Noise Separation, Test-Retest Reliability, and Group Discrimination.,” Neuroimage, vol. 117, pp. 67–79, Aug. 2015, doi: 10.1016/j.neuroimage.2015.05.015.

[11] S. Noble, D. Scheinost, and R. T. Constable, “A decade of test-retest reliability of functional connectivity: A systematic review and meta-analysis.,” Neuroimage, vol. 203, p. 116157, Dec. 2019, doi: 10.1016/j.neuroimage.2019.116157.

[12] E. Amico and J. Goñi, “The quest for identifiability in human functional connectomes,” Sci. Rep., vol. 8, no. 1, p. 8254, 2018, doi: 10.1038/s41598-018-25089-1.

[13] E. S. Finn et al., “Functional connectome fingerprinting: identifying individuals using patterns of brain connectivity,” Nat. Neurosci., vol. 18, no. 11, pp. 1664–1671, 2015, doi: 10.1038/nn.4135.

[14] X.-N. Zuo, C. Kelly, J. S. Adelstein, D. F. Klein, F. X. Castellanos, and M. P. Milham, “Reliable intrinsic connectivity networks: Test–retest evaluation using ICA and dual regression approach,” Neuroimage, vol. 49, no. 3, pp. 2163–2177, 2010, doi: 10.1016/j.neuroimage.2009.10.080.

[15] Y. Wang et al., “Intra-session test-retest reliability of functional connectivity in infants.,” Neuroimage, vol. 239, p. 118284, Oct. 2021, doi: 10.1016/j.neuroimage.2021.118284.

[16] P. E. Shrout and J. L. Fleiss, “Intraclass correlations: Uses in assessing rater reliability.,” 1979, American Psychological Association, US. doi: 10.1037/0033-2909.86.2.420.

[17] S. Noble, M. N. Spann, F. Tokoglu, X. Shen, R. T. Constable, and D. Scheinost, “Influences on the Test-Retest Reliability of Functional Connectivity MRI and its Relationship with Behavioral Utility.,” Cereb. Cortex, vol. 27, no. 11, pp. 5415–5429, Nov. 2017, doi: 10.1093/cercor/bhx230.

[18] L. M. Shah, J. A. Cramer, M. A. Ferguson, R. M. Birn, and J. S. Anderson, “Reliability and reproducibility of individual differences in functional connectivity acquired during task and resting state.,” Brain Behav., vol. 6, no. 5, p. e00456, May 2016, doi: 10.1002/brb3.456.

[19] Maxwell L Elliott et al., “What Is the Test-Retest Reliability of Common Task-Functional MRI Measures? New Empirical Evidence and a Meta-Analysis,” Psychol. Sci., vol. 31, no. 7, pp. 792–806, Jun. 2020, doi: 10.1177/0956797620916786.

[20] S. Noble, D. Scheinost, and R. T. Constable, “A guide to the measurement and interpretation of fMRI test-retest reliability.,” Curr. Opin. Behav. Sci., vol. 40, pp. 27–32, Aug. 2021, doi: 10.1016/j.cobeha.2020.12.012.

[21] C. C. Camp, S. Noble, D. Scheinost, A. Stringaris, and D. M. Nielson, “Test-Retest Reliability of Functional Connectivity in Adolescents With Depression.,” Biol. psychiatry. Cogn. Neurosci. neuroimaging, vol. 9, no. 1, pp. 21–29, Jan. 2024, doi: 10.1016/j.bpsc.2023.09.002.

[22] S. M. Smith et al., “Correspondence of the brain’s functional architecture during activation and rest.,” Proc. Natl. Acad. Sci. U. S. A., vol. 106, no. 31, pp. 13040–13045, Aug. 2009, doi: 10.1073/pnas.0905267106.

[23] B. T. T. Yeo et al., “The organization of the human cerebral cortex estimated by intrinsic functional connectivity.,” J. Neurophysiol., vol. 106, no. 3, pp. 1125–1165, Sep. 2011, doi: 10.1152/jn.00338.2011.

[24] A. B. Poppe, K. Wisner, G. Atluri, K. O. Lim, V. Kumar, and A. W. 3rd Macdonald, “Toward a neurometric foundation for probabilistic independent component analysis of fMRI data.,” Cogn. Affect. Behav. Neurosci., vol. 13, no. 3, pp. 641–659, Sep. 2013, doi: 10.3758/s13415-013-0180-8.

[25] K. R. A. Van Dijk, T. Hedden, A. Venkataraman, K. C. Evans, S. W. Lazar, and R. L. Buckner, “Intrinsic functional connectivity as a tool for human connectomics: theory, properties, and optimization.,” J. Neurophysiol., vol. 103, no. 1, pp. 297–321, Jan. 2010, doi: 10.1152/jn.00783.2009.

[26] R. M. Birn et al., “The effect of scan length on the reliability of resting-state fMRI connectivity estimates.,” Neuroimage, vol. 83, pp. 550–558, Dec. 2013, doi: 10.1016/j.neuroimage.2013.05.099.

[27] C. T. Whitlow, R. Casanova, and J. A. Maldjian, “Effect of resting-state functional MR imaging duration on stability of graph theory metrics of brain network connectivity.,” Radiology, vol. 259, no. 2, pp. 516–524, May 2011, doi: 10.1148/radiol.11101708.

[28] X.-H. Liao et al., “Functional brain hubs and their test-retest reliability: a multiband resting-state functional MRI study.,” Neuroimage, vol. 83, pp. 969–982, Dec. 2013, doi: 10.1016/j.neuroimage.2013.07.058.

[29] T. O. Laumann et al., “Functional System and Areal Organization of a Highly Sampled Individual Human Brain.,” Neuron, vol. 87, no. 3, pp. 657–670, Aug. 2015, doi: 10.1016/j.neuron.2015.06.037.

[30] E. M. Gordon et al., “Precision Functional Mapping of Individual Human Brains.,” Neuron, vol. 95, no. 4, pp. 791-807.e7, Aug. 2017, doi: 10.1016/j.neuron.2017.07.011.

[31] J. W. Cho, A. Korchmaros, J. T. Vogelstein, M. P. Milham, and T. Xu, “Impact of concatenating fMRI data on reliability for functional connectomics,” Neuroimage, vol. 226, p. 117549, 2021, doi: 10.1016/j.neuroimage.2020.117549.

[32] K. S. Button et al., “Power failure: why small sample size undermines the reliability of neuroscience,” Nat. Rev. Neurosci., vol. 14, no. 5, pp. 365–376, 2013, doi: 10.1038/nrn3475.

[33] S. Marek et al., “Reproducible brain-wide association studies require thousands of individuals,” Nature, vol. 603, no. 7902, pp. 654–660, 2022, doi: 10.1038/s41586-022-04492-9.

[34] L. Ma et al., “Effect of scanning duration and sample size on reliability in resting state fMRI dynamic causal modeling analysis,” Neuroimage, vol. 292, p. 120604, 2024, doi: 10.1016/j.neuroimage.2024.120604.

[35] B. O. Turner, E. J. Paul, M. B. Miller, and A. K. Barbey, “Small sample sizes reduce the replicability of task-based fMRI studies,” Commun. Biol., vol. 1, no. 1, p. 62, 2018, doi: 10.1038/s42003-018-0073-z.

[36] D. Szucs and J. P. A. Ioannidis, “Sample size evolution in neuroimaging research: An evaluation of highly-cited studies (1990–2012) and of latest practices (2017–2018) in high-impact journals,” Neuroimage, vol. 221, p. 117164, 2020, doi: 10.1016/j.neuroimage.2020.117164.

[37] R. A. Poldrack and M. J. Farah, “Progress and challenges in probing the human brain.,” Nature, vol. 526, no. 7573, pp. 371–379, Oct. 2015, doi: 10.1038/nature15692.

[38] B. Cassone et al., “TR(Acking) Individuals Down: Exploring the Effect of Temporal Resolution in Resting-State Functional MRI Fingerprinting,” Hum. Brain Mapp., vol. 46, no. 2, p. e70125, Feb. 2025, doi: 10.1002/hbm.70125.

[39] M.-S. Cahart et al., “Comparing the test–retest reliability of resting-state functional magnetic resonance imaging metrics across single band and multiband acquisitions in the context of healthy aging,” Hum. Brain Mapp., vol. 44, no. 5, pp. 1901–1912, Apr. 2023, doi: 10.1002/hbm.26180.

[40] J. Wang, J. Han, V. T. Nguyen, L. Guo, and C. C. Guo, “Improving the Test-Retest Reliability of Resting State fMRI by Removing the Impact of Sleep.,” Front. Neurosci., vol. 11, p. 249, 2017, doi: 10.3389/fnins.2017.00249.

[41] R. A. Poldrack et al., “Scanning the horizon: towards transparent and reproducible neuroimaging research,” Nat. Rev. Neurosci., vol. 18, no. 2, pp. 115–126, 2017, doi: 10.1038/nrn.2016.167.

[42] R. Patriat et al., “The effect of resting condition on resting-state fMRI reliability and consistency: A comparison between resting with eyes open, closed, and fixated,” Neuroimage, vol. 78, pp. 463–473, 2013, doi: 10.1016/j.neuroimage.2013.04.013.

[43] D. C. Van Essen, S. M. Smith, D. M. Barch, T. E. J. Behrens, E. Yacoub, and K. Ugurbil, “The WU-Minn Human Connectome Project: An overview,” Neuroimage, vol. 80, pp. 62–79, 2013, doi: 10.1016/j.neuroimage.2013.05.041.

[44] D. C. Van Essen et al., “The Human Connectome Project: a data acquisition perspective.,” Neuroimage, vol. 62, no. 4, pp. 2222–2231, Oct. 2012, doi: 10.1016/j.neuroimage.2012.02.018.

[45] S. M. Smith et al., “Resting-state fMRI in the Human Connectome Project,” Neuroimage, vol. 80, pp. 144–168, 2013, doi: 10.1016/j.neuroimage.2013.05.039.

[46] M. F. Glasser et al., “The minimal preprocessing pipelines for the Human Connectome Project.,” Neuroimage, vol. 80, pp. 105–124, Oct. 2013, doi: 10.1016/j.neuroimage.2013.04.127.

[47] M. Jenkinson, C. F. Beckmann, T. E. J. Behrens, M. W. Woolrich, and S. M. Smith, “FSL,” Neuroimage, vol. 62, no. 2, pp. 782–790, 2012, doi: 10.1016/j.neuroimage.2011.09.015.

[48] J. Bijsterbosch, S. Smith, and C. Beckmann, Introductin to Resting State fMRI Functional Connectivity, 1st ed. Oxford University Press, 2017.

[49] K. Somandepalli et al., “Short-term test-retest reliability of resting state fMRI metrics in children with and without attention-deficit/hyperactivity disorder.,” Dev. Cogn. Neurosci., vol. 15, pp. 83–93, Oct. 2015, doi: 10.1016/j.dcn.2015.08.003.

[50] A. S. Choe et al., “Reproducibility and Temporal Structure in Weekly Resting-State fMRI over a Period of 3.5 Years.,” PLoS One, vol. 10, no. 10, p. e0140134, 2015, doi: 10.1371/journal.pone.0140134.

[51] D. V Cicchetti and S. A. Sparrow, “Developing criteria for establishing interrater reliability of specific items: applications to assessment of adaptive behavior.,” Am. J. Ment. Defic., vol. 86, no. 2, pp. 127–137, Sep. 1981.

[52] L. Geerligs Cam-CAN, and R. N. Henson, “Functional connectivity and structural covariance between regions of interest can be measured more accurately using multivariate distance correlation,” Neuroimage, vol. 135, pp. 16–31, 2016, doi: 10.1016/j.neuroimage.2016.04.047.

[53] L. Geerligs, K. A. Tsvetanov Cam-Can, and R. N. Henson, “Challenges in measuring individual differences in functional connectivity using fMRI: The case of healthy aging.,” Hum. Brain Mapp., vol. 38, no. 8, pp. 4125–4156, Aug. 2017, doi: 10.1002/hbm.23653.

[54] E. S. Finn, D. Scheinost, D. M. Finn, X. Shen, X. Papademetris, and R. T. Constable, “Can brain state be manipulated to emphasize individual differences in functional connectivity?,” Neuroimage, vol. 160, pp. 140–151, 2017, doi: 10.1016/j.neuroimage.2017.03.064.

[55] L. Q. Uddin et al., “Controversies and progress on standardization of large-scale brain network nomenclature.,” Netw. Neurosci. (Cambridge, Mass.), vol. 7, no. 3, pp. 864–905, 2023, doi: 10.1162/netn_a_00323.

[56] L. R. Dice, “Measures of the Amount of Ecologic Association Between Species,” Ecology, vol. 26, no. 3, pp. 297–302, Jun. 1945, doi: 10.2307/1932409.

[57] A. F. Mejia et al., “Improved estimation of subject-level functional connectivity using full and partial correlation with empirical Bayes shrinkage,” Neuroimage, vol. 172, pp. 478–491, 2018, doi: 10.1016/j.neuroimage.2018.01.029.

[58] S. Noble et al., “Multisite reliability of MR-based functional connectivity,” Neuroimage, vol. 146, pp. 959–970, 2017, doi: 10.1016/j.neuroimage.2016.10.020.

[59] D. G. Tomasi, E. Shokri-Kojori, and N. D. Volkow, “Temporal Evolution of Brain Functional Connectivity Metrics: Could 7 Min of Rest be Enough?,” Cereb. Cortex, vol. 27, no. 8, pp. 4153–4165, Aug. 2017, doi: 10.1093/cercor/bhw227.

[60] C. Gratton et al., “Defining Individual-Specific Functional Neuroanatomy for Precision Psychiatry,” Biol. Psychiatry, vol. 88, no. 1, pp. 28–39, 2020, doi: 10.1016/j.biopsych.2019.10.026.

[61] A. F. Mejia et al., “Improving reliability of subject-level resting-state fMRI parcellation with shrinkage estimators,” Neuroimage, vol. 112, pp. 14–29, 2015, doi: 10.1016/j.neuroimage.2015.02.042.

[62] E. S. Finn and M. D. Rosenberg, “Beyond fingerprinting: Choosing predictive connectomes over reliable connectomes,” Neuroimage, vol. 239, p. 118254, 2021, doi: 10.1016/j.neuroimage.2021.118254.

[63] Philip A Kragel, Xiaochun Han, Thomas E Kraynak, Peter J Gianaros, and Tor D Wager, “Functional MRI Can Be Highly Reliable, but It Depends on What You Measure: A Commentary on Elliott et al. (2020),” Psychol. Sci., vol. 32, no. 4, pp. 622–626, Mar. 2021, doi: 10.1177/0956797621989730.

[64] R. X. Rodriguez, S. Noble, C. C. Camp, and D. Scheinost, “Connectome caricatures: removing large-amplitude co-activation patterns in resting-state fMRI emphasizes individual differences.,” bioRxiv Prepr. Serv. Biol., Apr. 2024, doi: 10.1101/2024.04.08.588578.

[65] A. R. Franco, M. V Mannell, V. D. Calhoun, and A. R. Mayer, “Impact of analysis methods on the reproducibility and reliability of resting-state networks.,” Brain Connect., vol. 3, no. 4, pp. 363–374, 2013, doi: 10.1089/brain.2012.0134.

[66] Y. Ma and A. W. MacDonald III, “Impact of Independent Component Analysis Dimensionality on the Test-Retest Reliability of Resting-State Functional Connectivity.,” Brain Connect., vol. 11, no. 10, pp. 875–886, Dec. 2021, doi: 10.1089/brain.2020.0970.

[67] E. M. Gordon, T. O. Laumann, B. Adeyemo, J. F. Huckins, W. M. Kelley, and S. E. Petersen, “Generation and Evaluation of a Cortical Area Parcellation from Resting-State Correlations,” Cereb. Cortex, vol. 26, no. 1, pp. 288–303, Jan. 2016, doi: 10.1093/cercor/bhu239.

[68] X.-N. Zuo et al., “An open science resource for establishing reliability and reproducibility in functional connectomics,” Sci. Data, vol. 1, no. 1, p. 140049, 2014, doi: 10.1038/sdata.2014.49.

[69] X.-N. Zuo and X.-X. Xing, “Test-retest reliabilities of resting-state FMRI measurements in human brain functional connectomics: A systems neuroscience perspective,” Neurosci. Biobehav. Rev., vol. 45, pp. 100–118, 2014, doi: 10.1016/j.neubiorev.2014.05.009.

