## Supplemental Information for "Test-retest reliability of resting-state fMRI functional connectivity: impact of scan length and number of participants"

**SUPPLEMENTARY INFORMATION**

1. *VOXEL-BASED RSN ANALYSIS*

Regarding spatial similarity, which quantifies the overlap between ICA-derived networks and Yeo's canonical RSNs, scan length had no significant main effect at the group-level, indicating that as little as 300 volumes (3.6 minutes) of data were sufficient for reliable estimates (Figure S1-S2, Table S1A). At the subject-level, however, all post hoc pairwise comparisons between scan lengths were significant, with the full 1200 volumes (14.4 minutes) scan yielding the highest Dice coefficients. Furthermore, for spatial similarity, there was also a significant interaction between scan length and RSN (Table S1C). Specifically, for all RSNs except the VN and LN, the full scan length produced the highest Dice coefficients (Table S1E). For the VN and LN, no statistically significant improvements were observed beyond 900 volumes (10.8 minutes).

For cohort size at the group-level (Figure S1, Table S2), no significant differences were observed beyond 50 subjects when considering all RSNs together. A significant interaction between the number of participants and RSN was detected. When analysed per RSN, the minimum cohort size beyond which additional participants did not lead to statistically significant differences was: 10 participants for the SMN and VN; 20 for the FPN and DMN; 50 for the DAN and VAN; and 100 for the LN. At the subject-level (Figure S2, Table S2), no overall main effect of cohort size was observed, nor was there a significant interaction between RSN and number of participants.

For spatial similarity (Table S3), the acquisition run (REST1_LR, REST1_RL, REST2_LR, REST2_RL) had a significant effect at both levels, indicating that the match between ICA-derived networks and Yeo's templates varied depending on the run.

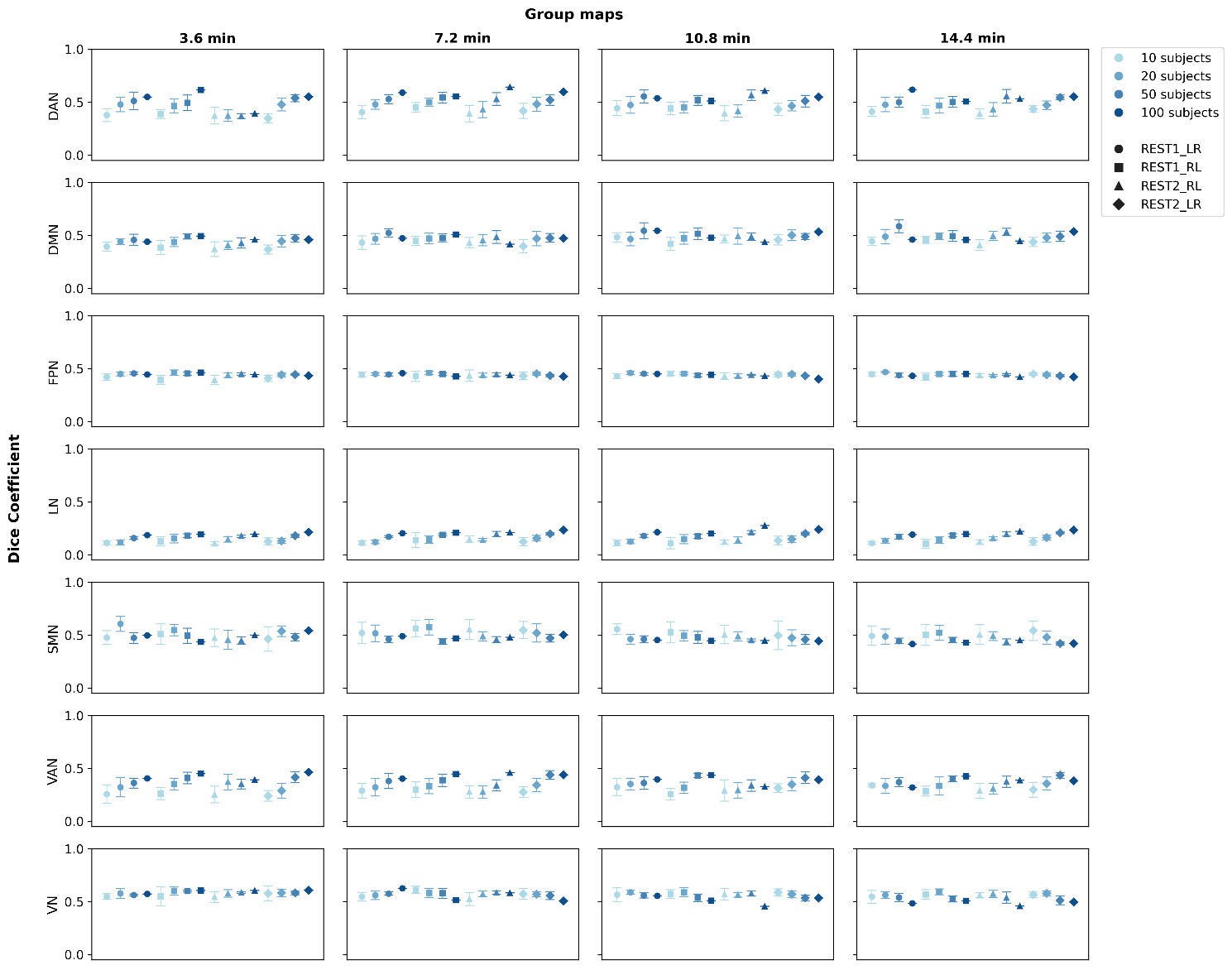

**Supplementary Figure S1 – Group-level spatial similarity of ICA-derived networks.** Mean Dice coefficient as a function of scan length (in minutes) and number of participants, averaged over 10 iterations for the full preprocessing pipeline. Error bars indicate standard deviation across iterations. Scan lengths correspond to 300 (3 minutes 36 seconds), 600 (7 minutes 12 seconds), 900 (10 minutes 48 seconds) and 1200 (14 minutes 24 seconds) volumes, respectively. Data points are slightly offset along the x-axis for visual clarity. DAN: dorsal attention network, DMN: default mode network, FPN: fronto-parietal network, LN: limbic network, SMN: somatomotor network, VAN: ventral attention network, VN: visual network.

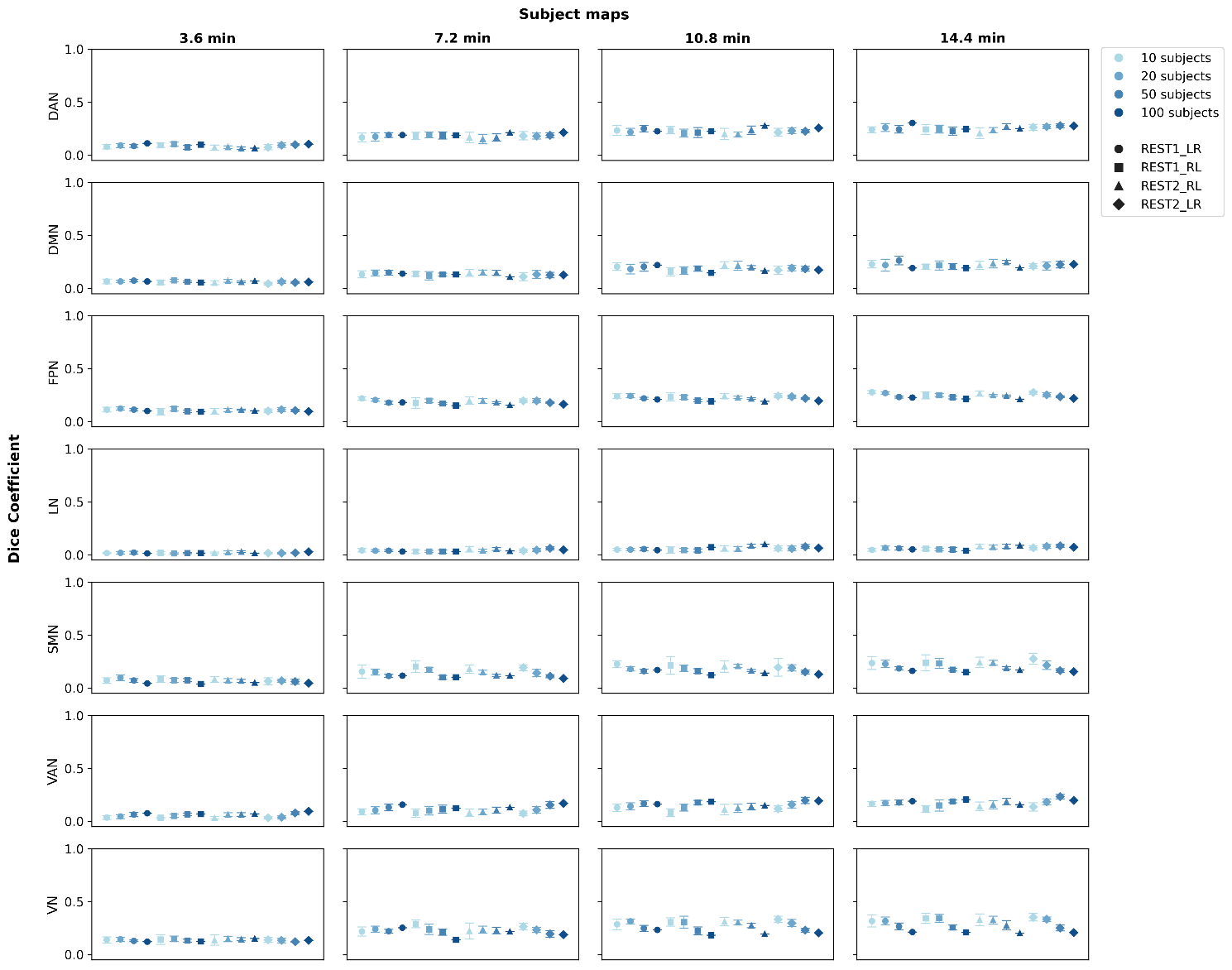

**Supplementary Figure S2 - Subject-level spatial similarity of ICA-derived networks.** Mean Dice coefficient as a function of scan length (in minutes) and number of participants, averaged over 10 iterations for full preprocessing pipeline. Error bars indicate standard deviation across iterations. Scan lengths correspond to 300 (3 minutes 36 seconds), 600 (7 minutes 12 seconds), 900 (10 minutes 48 seconds) and 1200 (14 minutes 24 seconds) volumes, respectively. Data points are slightly offset along the x-axis for visual clarity. DAN: dorsal attention network, DMN: default mode network, FPN: fronto-parietal network, LN: limbic network, SMN: somatomotor network, VAN: ventral attention network, VN: visual network.

2. *NODE-BASED CONNECTOME AND VOXEL-BASED RSN ANALYSIS*
   2.1. *Effects of Pipeline*

We also evaluated the impact of two preprocessing strategies on the reliability of resting-state measures. The preprocessing pipeline had a significant effect on all connectome-based metrics except fingerprint success rate (Figure S3, Table S6). Metrics consistently showed higher values when processed using the full pipeline compared to the minimal pipeline, suggesting that additional preprocessing steps, such as nuisance regression of motion parameters and WM and CSF mean time series, improved reliability across all metrics. The preprocessing pipeline had no statistically significant main effect on spatial reliability (Figure S4, Table S6).

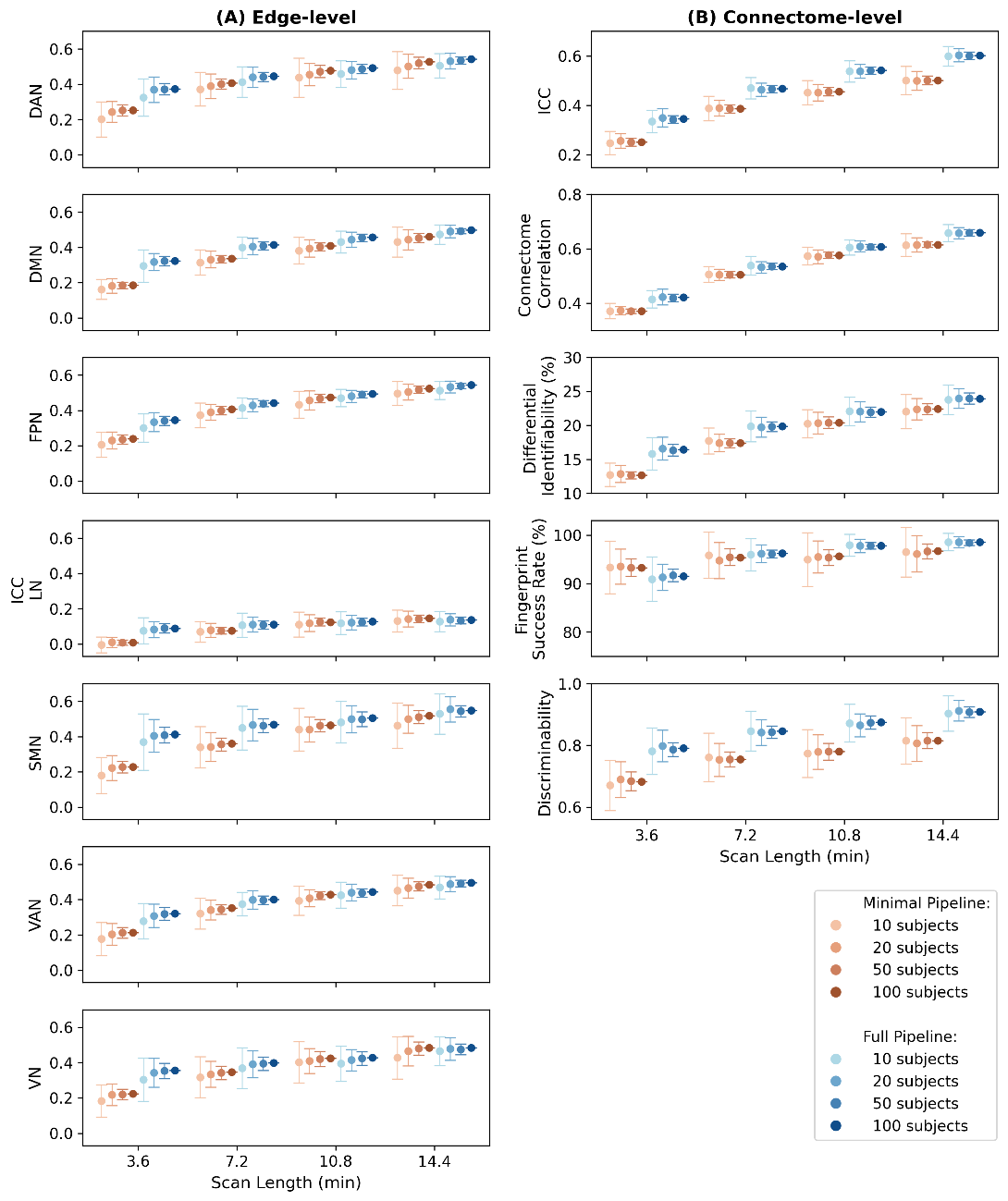

**Supplementary Figure S3 – Comparison between within-session minimal and full preprocessing pipelines for edge-level (A) and connectome-level (B) metrics, as a function of scan length (in minutes) and number of participants.** (A) Edge-level ICC per RSN. (B) Connectome-level ICC, connectome correlation, differential identifiability, fingerprinting success rate and discriminability. Dots represent the mean and error bars the standard deviation of the mean across 100 iterations (except for the case of 100 subjects, for which only 1 iteration is performed). Scan lengths correspond to 300 (3 minutes 36 seconds), 600 (7 minutes 12 seconds), 900 (10 minutes 48 seconds) and 1200 (14 minutes 24 seconds) volumes, respectively. Data points are slightly offset along the x-axis for visual clarity. DAN: dorsal attention network, DMN: default mode network, FPN: fronto-parietal network, LN: limbic network, SMN: somatomotor network, VAN: ventral attention network, VN: visual network.

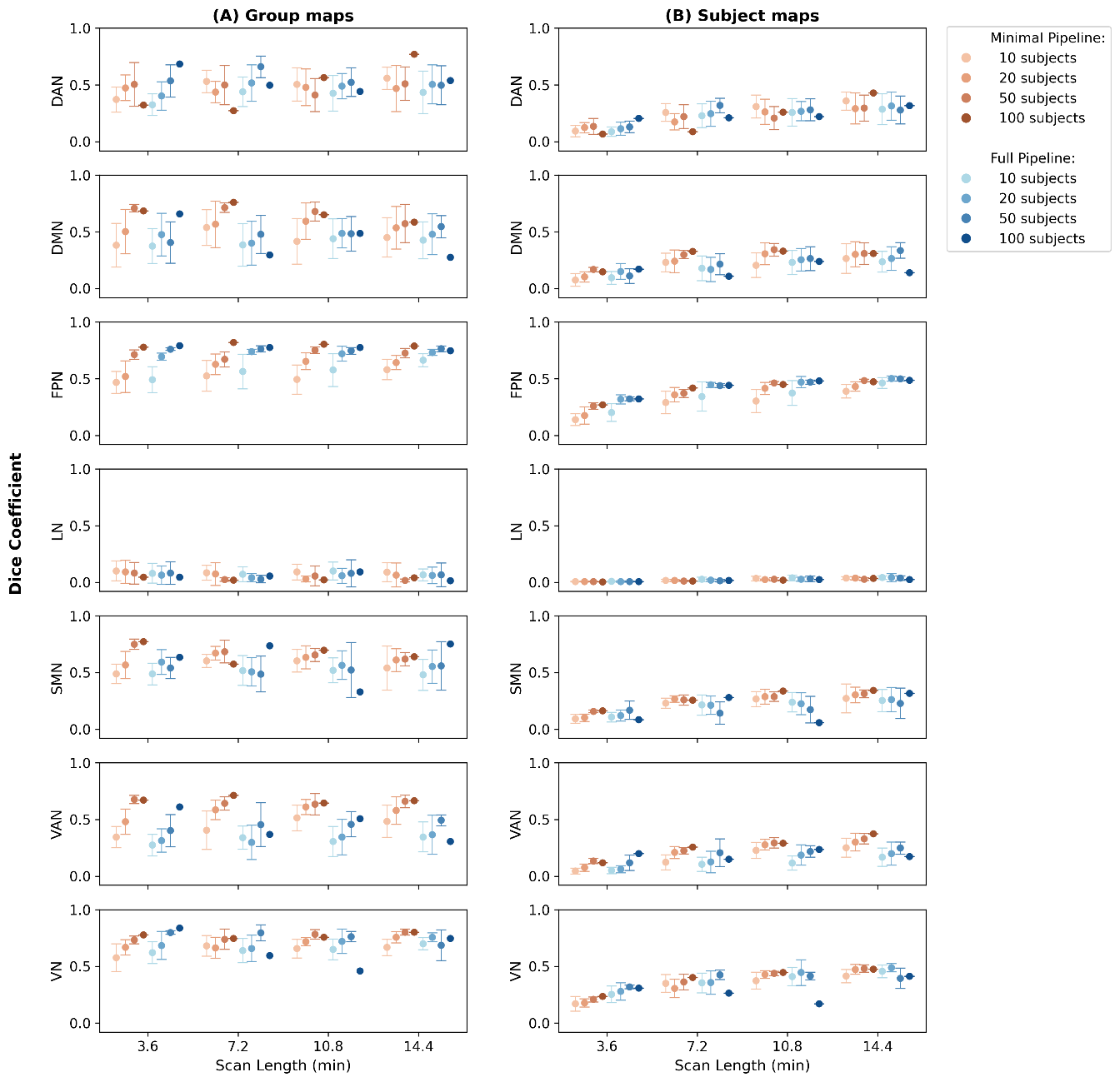

**Supplementary Figure S4 - Comparison between minimal and full preprocessing pipelines of within-session spatial test-retest reliability of ICA-derived networks.** Group-level (A) and subject-level (B) mean Dice coefficient as a function of scan duration in minutes and number of participants, averaged over 10 iterations. Error bars indicate standard deviation across iterations. Scan lengths correspond to 300 (3 minutes 36 seconds), 600 (7 minutes 12 seconds), 900 (10 minutes 48 seconds) and 1200 (14 minutes 24 seconds) volumes, respectively. Data points are slightly offset along the x-axis for visual clarity. DAN: dorsal attention network, DMN: default mode network, FPN: fronto-parietal network, LN: limbic network, SMN: somatomotor network, VAN: ventral attention network, VN: visual network.

2.3. *Effects of RSN size*

In order to evaluate the relationship between network size and each metric value, we correlated network size with each FC metric, averaging values across all sessions. For node-based metrics, network size was defined as the number of parcels from the Schaefer atlas corresponding to each RSN. For voxel-based metrics, network size was defined as the number of voxels in each RSN template.

A positive correlation was observed between RSN template size and mean Dice coefficient across all analyses. The LN, which had the smallest template (11 547 voxels), consistently exhibited the lowest Dice values across both metrics and analysis levels. In contrast, larger networks such as VN (22 282 voxels) and DMN (30 127 voxels) tended to show higher spatial overlap (Figure S5). This pattern suggests that Dice coefficients may be inherently sensitive to template size.

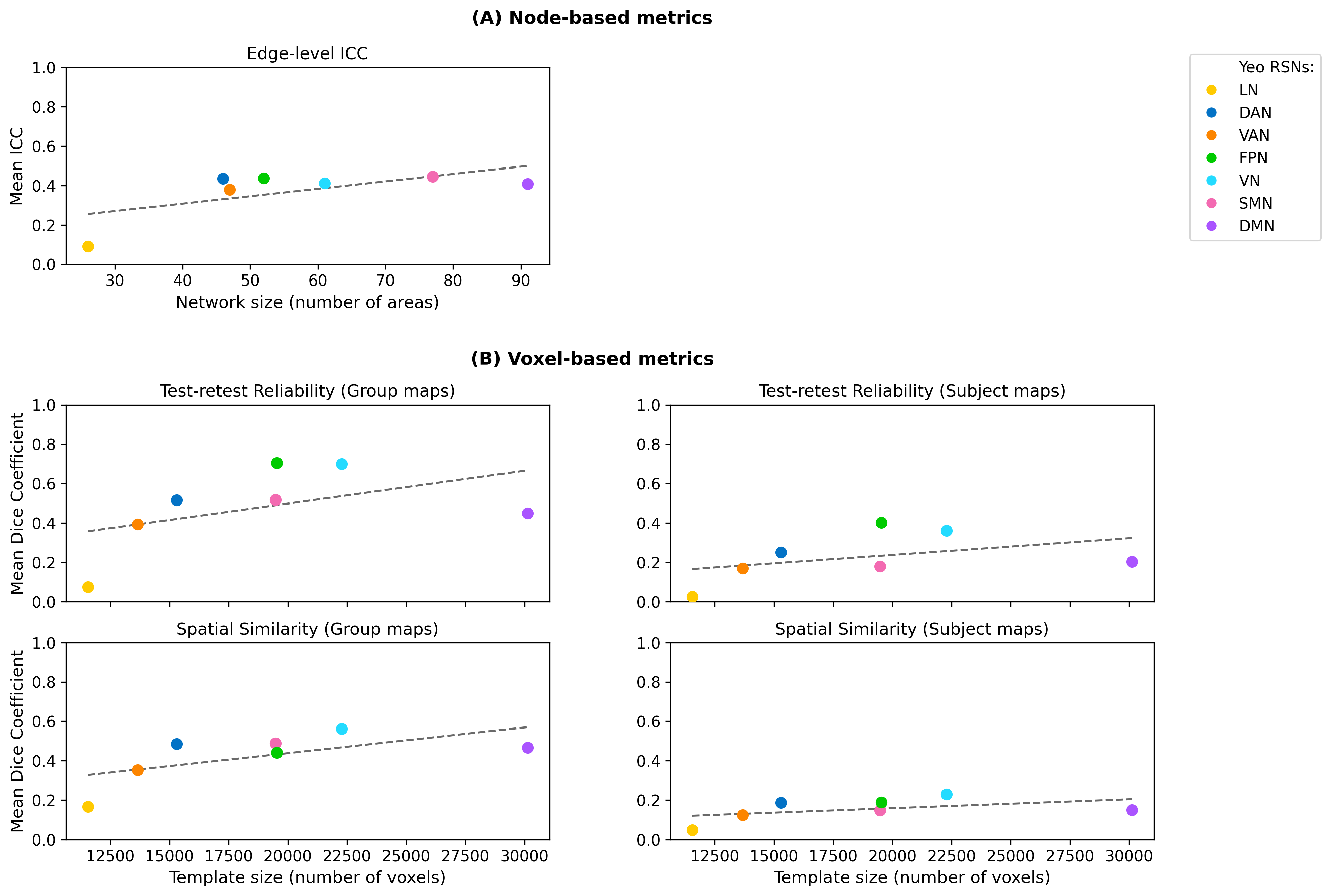

**Supplementary Figure S5 – Relationship between network size and reliability metrics.** (A) Node-based metrics: network size, defined as the number of nodes per RSN according to the 400-node Schaefer atlas, plotted against mean edge-level ICC averaged across all conditions (scan length, number of subjects, and session type). (B) Voxel-based metrics: template size, defined as the number of voxels in the binary mask of each Yeo RSN, plotted against the mean Dice coefficient for spatial test-retest reliability (between sessions and runs) and spatial similarity, averaged across all conditions (scan length, number of subjects, and session type). DAN: dorsal attention network; DMN: default mode network; FPN: fronto-parietal network; LN: limbic network; SMN: somatomotor network; VAN: ventral attention network; VN: visual network.

4. *STATISTICAL ANALYSIS*

Full statistics are provided in Supplementary Tables S1–S6, organized by the variable analyzed: S1 presents the effects of scan length on test–retest reliability of connectivity metrics; S2 presents the effects of number of subjects; S3 presents the effects of test–retest condition; S4 presents the effects of resting-state network; S5 presents the effects of temporal downsampling strategy; and S6 presents the effects of preprocessing pipeline. Each table presents results for all connectivity metrics. The statistical models used for each analysis are summarised below:

| **Analysis** | **Metric type** | **Model formula** |
| --- | --- | --- |
| Scan Length, Number of Subjects, Test-retest condition | All node-based metrics except ICC edge-level | *mean_value ~ scan_length * n_subs + session/run* |
| Scan Length, Number of Subjects, Test-retest condition, Resting-State Network | RSN-level metrics (ICC edge, voxel-based) | *mean_value ~ scan_length * n_subs * rsn + session/run* |
| Temporal Downsampling strategy | All node-based metrics except ICC edge-level | *mean_value ~ scan_length * strategy + session/run* |
| Temporal Downsampling strategy | RSN-level metrics (ICC edge, voxel-based) | *mean_value ~ scan_length * strategy * rsn + session/run* |
| Preprocessing pipeline | All node-based metrics except ICC edge-level | *mean_value ~ scan_length * n_subs * pipeline + session/run* |
| Preprocessing pipeline | RSN-level metrics (ICC edge, voxel-based) | *mean_value ~ scan_length * n_subs * pipeline * rsn + session/run* |

**Supplementary Table S1 – Effects of scan length on test–retest reliability of connectivity metrics. (A) Main effect of scan length (ANOVA).** ANOVA results for the effect of scan length on all node‑based and voxel‑based reliability metrics. For most metrics, the model was mean_value ~ scan_length * n_subs + session/run; for edge‑level ICC, test–retest reliability and spatial similarity, the model was mean_value ~ scan_length * n_subs * rsn + session/run. **(B)** **Pairwise comparisons across scan lengths.** Post‑hoc pairwise contrasts between scan lengths for each reliability metric corresponding to (A). **(C) Interaction between scan length and resting‑state network.** ANOVA results for the interaction between scan length and RSN on edge‑level ICC, test–retest reliability and spatial similarity. **(D) Effect of scan length within each resting‑state network.** ANOVA results for the effect of scan length computed separately for each RSN; in this case, the model was mean_value ~ scan_length * n_subs + session/run for each RSN. **(E) Pairwise scan length comparisons within each resting‑state network.** Post‑hoc pairwise contrasts between scan lengths for each RSN corresponding to (D). Significance levels (Bonferroni corrected for 10 comparison/metrics): *** p < 0.0001, ** p < 0.001, * p < 0.005.

| **(A) *Main effect of Scan Length (ANOVA)*** | | | | |
| --- | --- | --- | --- | --- |
| **Metric** | **Sum Sq** | **Df** | **F value** | ***p* value** |
| Edge-level ICC | 0.039340 | 3 | 31.7216 | 6.928e-15 *** |
| Connectome-level ICC | 0.075804 | 3 | 1114.1600 | < 2.2e-16 *** |
| Connectome correlation | 0.06421 | 3 | 2212.2673 | < 2.2e-16 *** |
| Differential Identifiability | 0.006552 | 3 | 251.0284 | 4.865e-13 *** |
| Success rate | 59.0 | 3 | 81.9210 | 1.579e-09 *** |
| Discriminability | 0.01568 | 3 | 102.7226 | 3.174e-10 *** |
| Test-Retest Reliability (group maps) | 0.01538 | 3 | 1.2407 | 0.298456 |
| Test-Retest Reliability (subject maps) | 0.038591 | 3 | 9.5387 | 1.172e-05 *** |
| Spatial Similarity (group maps) | 0.00738 | 3 | 3.4921 | 0.0159571 |
| Spatial Similarity (subject maps) | 0.061182 | 3 | 91.7011 | < 2.2e-16 *** |

| **(B) *Pairwise comparisons across Scan Lengths*** | | | | | |
| --- | --- | --- | --- | --- | --- |
| ***Edge-level ICC*** | | | | | |
| **Contrast** | **Estimate** | **SE** | **df** | **t.ratio** | **p.value** |
| scan_length300 − scan_length600 | -0.0738 | 0.00384 | 111 | -19.199 | <0.0001 *** |
| scan_length300 − scan_length900 | -0.1107 | 0.00384 | 111 | -28.804 | <0.0001 *** |
| scan_length300 − scan_length1200 | -0.1458 | 0.00384 | 111 | -37.942 | <0.0001 *** |
| scan_length600 − scan_length900 | -0.0369 | 0.00384 | 111 | -9.605 | <0.0001 *** |
| scan_length600 − scan_length1200 | -0.0720 | 0.00384 | 111 | -18.743 | <0.0001 *** |
| scan_length900 − scan_length1200 | -0.0351 | 0.00384 | 111 | -9.138 | <0.0001 *** |
| ***Connectome-level ICC*** | | | | | |
| **Contrast** | **Estimate** | **SE** | **df** | **t.ratio** | **p.value** |
| scan_length300 − scan_length600 | -0.1251 | 0.00238 | 15 | -52.549 | <0.0001 *** |
| scan_length300 − scan_length900 | -0.2007 | 0.00238 | 15 | -84.277 | <0.0001 *** |
| scan_length300 − scan_length1200 | -0.2556 | 0.00238 | 15 | -107.355 | <0.0001 *** |
| scan_length600 − scan_length900 | -0.0756 | 0.00238 | 15 | -31.729 | <0.0001 *** |
| scan_length600 − scan_length1200 | -0.1305 | 0.00238 | 15 | -54.806 | <0.0001 *** |
| scan_length900 − scan_length1200 | -0.0549 | 0.00238 | 15 | -23.077 | <0.0001 *** |
| ***Connectome correlation*** | | | | | |
| **Contrast** | **Estimate** | **SE** | **df** | **t.ratio** | **p.value** |
| scan_length300 − scan_length600 | -0.1153 | 0.00156 | 15 | -74.170 | <0.0001 *** |
| scan_length300 − scan_length900 | -0.1873 | 0.00156 | 15 | -120.465 | <0.0001 *** |
| scan_length300 − scan_length1200 | -0.2356 | 0.00156 | 15 | -151.466 | <0.0001 *** |
| scan_length600 − scan_length900 | -0.0720 | 0.00156 | 15 | -46.296 | <0.0001 *** |
| scan_length600 − scan_length1200 | -0.1202 | 0.00156 | 15 | -77.296 | <0.0001 *** |
| scan_length900 − scan_length1200 | -0.0482 | 0.00156 | 15 | -31.000 | <0.0001 *** |
| ***Differential Identifiability*** | | | | | |
| **Contrast** | **Estimate** | **SE** | **df** | **t.ratio** | **p.value** |
| scan_length300 − scan_length600 | -0.0358 | 0.00147 | 15 | -24.301 | <0.0001 *** |
| scan_length300 − scan_length900 | -0.0563 | 0.00147 | 15 | -38.167 | <0.0001 *** |
| scan_length300 − scan_length1200 | -0.0726 | 0.00147 | 15 | -49.211 | <0.0001 *** |
| scan_length600 − scan_length900 | -0.0204 | 0.00147 | 15 | -13.867 | <0.0001 *** |
| scan_length600 − scan_length1200 | -0.0367 | 0.00147 | 15 | -24.911 | <0.0001 *** |
| scan_length900 − scan_length1200 | -0.0163 | 0.00147 | 15 | -11.044 | <0.0001 *** |
| ***Success rate*** | | | | | |
| **Contrast** | **Estimate** | **SE** | **df** | **t.ratio** | **p.value** |
| scan_length300 − scan_length600 | -4.478 | 0.245 | 15 | -18.285 | <0.0001 *** |
| scan_length300 − scan_length900 | -5.909 | 0.245 | 15 | -24.126 | <0.0001 *** |
| scan_length300 − scan_length1200 | -6.349 | 0.245 | 15 | -25.925 | <0.0001 *** |
| scan_length600 − scan_length900 | -1.430 | 0.245 | 15 | -5.840 | 0.0002 ** |
| scan_length600 − scan_length1200 | -1.871 | 0.245 | 15 | -7.640 | <0.0001 *** |
| scan_length900 − scan_length1200 | -0.441 | 0.245 | 15 | -1.799 | 0.5526 |
| ***Discriminability*** | | | | | |
| **Contrast** | **Estimate** | **SE** | **df** | **t.ratio** | **p.value** |
| scan_length300 − scan_length600 | -0.0577 | 0.00357 | 15 | -16.176 | <0.0001 *** |
| scan_length300 − scan_length900 | -0.0874 | 0.00357 | 15 | -24.495 | <0.0001 *** |
| scan_length300 − scan_length1200 | -0.1147 | 0.00357 | 15 | -32.152 | <0.0001 *** |
| scan_length600 − scan_length900 | -0.0297 | 0.00357 | 15 | -8.319 | <0.0001 *** |
| scan_length600 − scan_length1200 | -0.0570 | 0.00357 | 15 | -15.976 | <0.0001 *** |
| scan_length900 − scan_length1200 | -0.0273 | 0.00357 | 15 | -7.657 | <0.0001 *** |
| ***Test-Retest Reliability (subject maps)*** | | | | | |
| **Contrast** | **Estimate** | **SE** | **df** | **t.ratio** | **p.value** |
| scan_length300 − scan_length600 | -0.0726 | 0.00694 | 111 | -10.464 | <0.0001 *** |
| scan_length300 − scan_length900 | -0.0999 | 0.00694 | 111 | -14.392 | <0.0001 *** |
| scan_length300 − scan_length1200 | -0.1346 | 0.00694 | 111 | -19.394 | <0.0001 *** |
| scan_length600 − scan_length900 | -0.0273 | 0.00694 | 111 | -3.928 | 0.0009 ** |
| scan_length600 − scan_length1200 | -0.0620 | 0.00694 | 111 | -8.930 | <0.0001 *** |
| scan_length900 − scan_length1200 | -0.0347 | 0.00694 | 111 | -5.002 | <0.0001 *** |
| ***Spatial Similarity (subject maps)*** | | | | | |
| **Contrast** | **Estimate** | **SE** | **df** | **t.ratio** | **p.value** |
| scan_length300 − scan_length600 | -0.0693 | 0.00199 | 333 | -34.772 | <0.0001 *** |
| scan_length300 − scan_length900 | -0.1074 | 0.00199 | 333 | -53.912 | <0.0001 *** |
| scan_length300 − scan_length1200 | -0.1298 | 0.00199 | 333 | -65.128 | <0.0001 *** |
| scan_length600 − scan_length900 | -0.0381 | 0.00199 | 333 | -19.140 | <0.0001 *** |
| scan_length600 − scan_length1200 | -0.0605 | 0.00199 | 333 | -30.355 | <0.0001 *** |
| scan_length900 − scan_length1200 | -0.0224 | 0.00199 | 333 | -11.216 | <0.0001 *** |

| **(C) *Interaction between Scan Length and Resting-State Network (ANOVA)*** | | | | |
| --- | --- | --- | --- | --- |
| **Metric** | **Sum Sq** | **Df** | **F value** | ***p* value** |
| Edge-level ICC | 0.023064 | 18 | 3.0996 | 0.0001357 ** |
| Test-Retest Reliability (group maps) | 0.02988 | 18 | 0.4019 | 0.985157 |
| Test-Retest Reliability (subject maps) | 0.032818 | 18 | 1.3519 | 0.1706784 |
| Spatial Similarity (group maps) | 0.01369 | 18 | 1.0795 | 0.3714577 |
| Spatial Similarity (subject maps) | 0.040123 | 18 | 10.0229 | < 2.2e-16 *** |

| **(D) *Effect of Scan Length within each Resting‑State Network (ANOVA)*** | | | | | |
| --- | --- | --- | --- | --- | --- |
| **Metric** | **RSN** | **Sum Sq** | **Df** | **F value** | ***p* value** |
| Spatial Similarity  (subject maps) | DAN | 0.061182 | 3 | 78.1709 | < 2.2e-16 *** |
|  | DMN | 0.060533 | 3 | 101.5916 | < 2.2e-16 *** |
|  | FPN | 0.062865 | 3 | 515.3891 | < 2.2e-16 *** |
|  | LN | 0.0045982 | 3 | 18.7747 | 4.832e-08 *** |
|  | SMN | 0.066393 | 3 | 140.5566 | < 2.2e-16 *** |
|  | VAN | 0.0240036 | 3 | 40.0738 | 9.138e-13 *** |
|  | VN | 0.091179 | 3 | 77.3668 | < 2.2e-16 *** |

| ***(E) Pairwise Scan Length comparisons within each Resting‑State Network*** | | | | | | | | |
| --- | --- | --- | --- | --- | --- | --- | --- | --- |
| **Metric** | **RSN** | **Contrast** | **Estimate** | | **SE** | **df** | **t.ratio** | **p.value** |
| Spatial Similarity  (subject  maps) | DAN | scan_length300 − scan_length600 | | -0.0968 | 0.00571 | 45 | -16.942 | <0.0001 *** |
|  |  | scan_length300 − scan_length900 | | -0.1417 | 0.00571 | 45 | -24.812 | <0.0001 *** |
|  |  | scan_length300 − scan_length1200 | | -0.1673 | 0.00571 | 45 | -29.303 | <0.0001 *** |
|  |  | scan_length600 − scan_length900 | | -0.0449 | 0.00571 | 45 | -7.870 | <0.0001 *** |
|  |  | scan_length600 − scan_length1200 | | -0.0706 | 0.00571 | 45 | -12.361 | <0.0001 *** |
|  |  | scan_length900 − scan_length1200 | | -0.0256 | 0.00571 | 45 | -4.491 | 0.0003 ** |
|  | DMN | scan_length300 − scan_length600 | | -0.0709 | 0.00498 | 45 | -14.221 | <0.0001 *** |
|  |  | scan_length300 − scan_length900 | | -0.1249 | 0.00498 | 45 | -25.072 | <0.0001 *** |
|  |  | scan_length300 − scan_length1200 | | -0.1561 | 0.00498 | 45 | -31.338 | <0.0001 *** |
|  |  | scan_length600 − scan_length900 | | -0.0541 | 0.00498 | 45 | -10.851 | <0.0001 *** |
|  |  | scan_length600 − scan_length1200 | | -0.0853 | 0.00498 | 45 | -17.116 | <0.0001 *** |
|  |  | scan_length900 − scan_length1200 | | -0.0312 | 0.00498 | 45 | -6.266 | <0.0001 *** |
|  | FPN | scan_length300 − scan_length600 | | -0.0785 | 0.00225 | 45 | -34.812 | <0.0001 *** |
|  |  | scan_length300 − scan_length900 | | -0.1156 | 0.00225 | 45 | -51.267 | <0.0001 *** |
|  |  | scan_length300 − scan_length1200 | | -0.1380 | 0.00225 | 45 | -61.221 | <0.0001 *** |
|  |  | scan_length600 − scan_length900 | | -0.0371 | 0.00225 | 45 | -16.454 | <0.0001 *** |
|  |  | scan_length600 − scan_length1200 | | -0.0595 | 0.00225 | 45 | -26.409 | <0.0001 *** |
|  |  | scan_length900 − scan_length1200 | | -0.0224 | 0.00225 | 45 | -9.954 | <0.0001 *** |
|  | LN | scan_length300 − scan_length600 | | -0.0220 | 0.00319 | 45 | -6.887 | <0.0001 *** |
|  |  | scan_length300 − scan_length900 | | -0.0415 | 0.00319 | 45 | -12.978 | <0.0001 *** |
|  |  | scan_length300 − scan_length1200 | | -0.0466 | 0.00319 | 45 | -14.573 | <0.0001 *** |
|  |  | scan_length600 − scan_length900 | | -0.0195 | 0.00319 | 45 | -6.091 | <0.0001 *** |
|  |  | scan_length600 − scan_length1200 | | -0.0246 | 0.00319 | 45 | -7.686 | <0.0001 *** |
|  |  | scan_length900 − scan_length1200 | | -0.0051 | 0.00319 | 45 | -1.596 | 0.7055 |
|  | SMN | scan_length300 − scan_length600 | | -0.0726 | 0.00444 | 45 | -16.371 | <0.0001 *** |
|  |  | scan_length300 − scan_length900 | | -0.1091 | 0.00444 | 45 | -24.582 | <0.0001 *** |
|  |  | scan_length300 − scan_length1200 | | -0.1379 | 0.00444 | 45 | -31.087 | <0.0001 *** |
|  |  | scan_length600 − scan_length900 | | -0.0364 | 0.00444 | 45 | -8.211 | <0.0001 *** |
|  |  | scan_length600 − scan_length1200 | | -0.0653 | 0.00444 | 45 | -14.716 | <0.0001 *** |
|  |  | scan_length900 − scan_length1200 | | -0.0289 | 0.00444 | 45 | -6.505 | <0.0001 *** |
|  | VAN | scan_length300 − scan_length600 | | -0.0560 | 0.005 | 45 | -11.213 | <0.0001 *** |
|  |  | scan_length300 − scan_length900 | | -0.0905 | 0.005 | 45 | -18.116 | <0.0001 *** |
|  |  | scan_length300 − scan_length1200 | | -0.1151 | 0.005 | 45 | -23.038 | <0.0001 *** |
|  |  | scan_length600 − scan_length900 | | -0.0345 | 0.005 | 45 | -6.903 | <0.0001 *** |
|  |  | scan_length600 − scan_length1200 | | -0.0591 | 0.005 | 45 | -11.825 | <0.0001 *** |
|  |  | scan_length900 − scan_length1200 | | -0.0246 | 0.005 | 45 | -4.922 | <0.0001 *** |
|  | VN | scan_length300 − scan_length600 | | -0.0883 | 0.00701 | 45 | -12.605 | <0.0001 *** |
|  |  | scan_length300 − scan_length900 | | -0.1289 | 0.00701 | 45 | -18.388 | <0.0001 *** |
|  |  | scan_length300 − scan_length1200 | | -0.1475 | 0.00701 | 45 | -21.044 | <0.0001 *** |
|  |  | scan_length600 − scan_length900 | | -0.0405 | 0.00701 | 45 | -5.783 | <0.0001 *** |
|  |  | scan_length600 − scan_length1200 | | -0.0591 | 0.00701 | 45 | -8.439 | <0.0001 *** |
|  |  | scan_length900 − scan_length1200 | | -0.0186 | 0.00701 | 45 | -2.656 | 0.0655 |

**Supplementary Table S2 – Effects of number of subjects on test–retest reliability of connectivity metrics. (A) Main effect of number of subjects (ANOVA).** ANOVA results for the effect of number of subjects on all node‑based and voxel‑based reliability metrics. For most metrics, the model was mean_value ~ scan_length * n_subs + session/run; for edge‑level ICC, test–retest reliability and spatial similarity, the model was mean_value ~ scan_length * n_subs * rsn + session/run. **(B) Pairwise comparisons across numbers of subjects.** Post‑hoc pairwise contrasts between numbers of subjects for each reliability metric corresponding to (A). **(C) Interaction between number of subjects and resting‑state network.** ANOVA results for the interaction between number of subjects and RSN on edge‑level ICC, test–retest reliability and spatial similarity. **(D) Effect of number of subjects within each resting‑state network**. ANOVA results for the effect of number of subjects computed separately for each RSN; in this case, the model was mean_value ~ scan_length * n_subs + session/run for each RSN. **(E) Pairwise number of subjects comparisons within each resting‑state network.** Post‑hoc pairwise contrasts between numbers of subjects for each RSN corresponding to (D). **(F) Interaction between scan length and number of subjects.** ANOVA results for the interaction between scan length and number of subjects on reliability metrics. Significance levels (Bonferroni corrected for 10 comparison/metrics): *** p < 0.0001, ** p < 0.001, * p < 0.005.

| **(A) *Main effect of Number of Subjects (ANOVA)*** | | | | |
| --- | --- | --- | --- | --- |
| **Metric** | **Sum Sq** | **Df** | **F value** | ***p* value** |
| Edge-level ICC | 0.002660 | 3 | 2.1451 | 0.0985935 |
| Connectome-level ICC | 0.000038 | 3 | 0.5618 | 0.6484 |
| Connectome correlation | 0.00003 | 3 | 1.0134 | 0.414251 |
| Differential Identifiability | 0.000030 | 3 | 1.1561 | 0.3590 |
| Success rate | 0.3 | 3 | 0.3987 | 0.755906 |
| Discriminability | 0.00008 | 3 | 0.5043 | 0.6851 |
| Test-Retest Reliability (group maps) | 0.14358 | 3 | 11.5867 | 1.152e-06 *** |
| Test-Retest Reliability (subject maps) | 0.009743 | 3 | 2.4083 | 0.0709251 |
| Spatial Similarity (group maps) | 0.05165 | 3 | 24.4460 | 2.543e-14 *** |
| Spatial Similarity (subject maps) | 0.000672 | 3 | 1.0068 | 0.38988 |

| **(B) *Pairwise comparisons across Number of Subjects*** | | | | | |
| --- | --- | --- | --- | --- | --- |
| ***Test-Retest Reliability (group maps)*** | | | | | |
| **Contrast** | **Estimate** | **SE** | **df** | **t.ratio** | **p.value** |
| n_subs10 − n_subs20 | -0.0485 | 0.0121 | 111 | -3.993 | 0.0007 ** |
| n_subs10 − n_subs50 | -0.1010 | 0.0121 | 111 | -8.315 | <0.0001 *** |
| n_subs10 − n_subs100 | -0.0907 | 0.0121 | 111 | -7.465 | <0.0001 *** |
| n_subs20 − n_subs50 | -0.0525 | 0.0121 | 111 | -4.323 | 0.0002 ** |
| n_subs20 − n_subs100 | -0.0422 | 0.0121 | 111 | -3.472 | 0.0044 * |
| n_subs50 − n_subs100 | 0.0103 | 0.0121 | 111 | 0.850 | 1.0000 |
| ***Spatial Similarity (group maps)*** | | | | | |
| **Contrast** | **Estimate** | **SE** | **df** | **t.ratio** | **p.value** |
| n_subs10 − n_subs20 | -0.02679 | 0.00355 | 333 | -7.554 | <0.0001 *** |
| n_subs10 − n_subs50 | -0.04402 | 0.00355 | 333 | -12.413 | <0.0001 *** |
| n_subs10 − n_subs100 | -0.05013 | 0.00355 | 333 | -14.134 | <0.0001 *** |
| n_subs20 − n_subs50 | -0.01723 | 0.00355 | 333 | -4.859 | <0.0001 *** |
| n_subs20 − n_subs100 | -0.02334 | 0.00355 | 333 | -6.581 | <0.0001 *** |
| n_subs50 − n_subs100 | -0.00611 | 0.00355 | 333 | -1.722 | 0.5165 |

| **(C) *Interaction between Number of Subjects and Resting-State Network (ANOVA)*** | | | | |
| --- | --- | --- | --- | --- |
| **Metric** | **Sum Sq** | **Df** | **F value** | ***p* value** |
| Edge-level ICC | 0.000935 | 18 | 0.1256 | 0.9999952 |
| Test-Retest Reliability (group maps) | 0.16592 | 18 | 2.2316 | 0.005807 |
| Test-Retest Reliability (subject maps) | 0.037448 | 18 | 1.5427 | 0.0886237 |
| Spatial Similarity (group maps) | 0.06378 | 18 | 5.0311 | 3.849e-10 *** |
| Spatial Similarity (subject maps) | 0.008336 | 18 | 2.0824 | 0.00634 |

| **(D) *Effect of Number of Subjects within each Resting‑State Network (ANOVA)*** | | | | | |
| --- | --- | --- | --- | --- | --- |
| **Metric** | **RSN** | **Sum Sq** | **Df** | **F value** | ***p* value** |
| Spatial Similarity  (group maps) | DAN | 0.05165 | 3 | 10.6389 | 2.084e-05 *** |
|  | DMN | 0.01899 | 3 | 9.0389 | 8.499e-05 *** |
|  | FPN | 0.00594 | 3 | 19.1408 | 3.806e-08 *** |
|  | LN | 0.015234 | 3 | 37.5964 | 2.553e-12 *** |
|  | SMN | 0.00936 | 3 | 4.5758 | 0.007016 |
|  | VAN | 0.068137 | 3 | 27.3557 | 3.179e-10 *** |
|  | VN | 0.00387 | 3 | 2.2466 | 0.0958200 |

| ***(E) Pairwise Number of Subjects comparisons within each Resting‑State Network*** | | | | | | | |
| --- | --- | --- | --- | --- | --- | --- | --- |
| **Metric** | **RSN** | **Contrast** | **Estimate** | **SE** | **df** | **t.ratio** | **p.value** |
| Spatial  Similarity  (group maps) | DAN | n_subs10 − n_subs20 | -0.0516 | 0.0142 | 45 | -3.628 | 0.0044 * |
|  |  | n_subs10 − n_subs50 | -0.1113 | 0.0142 | 45 | -7.827 | <0.0001 *** |
|  |  | n_subs10 − n_subs100 | -0.1502 | 0.0142 | 45 | -10.562 | <0.0001 *** |
|  |  | n_subs20 − n_subs50 | -0.0597 | 0.0142 | 45 | -4.199 | 0.0008 ** |
|  |  | n_subs20 − n_subs100 | -0.0986 | 0.0142 | 45 | -6.934 | <0.0001 *** |
|  |  | n_subs50 − n_subs100 | -0.0389 | 0.0142 | 45 | -2.735 | 0.0534 |
|  | DMN | n_subs10 − n_subs20 | -0.04242 | 0.00936 | 45 | -4.534 | 0.0003 ** |
|  |  | n_subs10 − n_subs50 | -0.07177 | 0.00936 | 45 | -7.671 | <0.0001 *** |
|  |  | n_subs10 − n_subs100 | -0.05110 | 0.00936 | 45 | -5.462 | <0.0001 *** |
|  |  | n_subs20 − n_subs50 | -0.02935 | 0.00936 | 45 | -3.136 | 0.0181 |
|  |  | n_subs20 − n_subs100 | -0.00868 | 0.00936 | 45 | -0.928 | 1.0000 |
|  |  | n_subs50 − n_subs100 | 0.02067 | 0.00936 | 45 | 2.209 | 0.1940 |
|  | FPN | n_subs10 − n_subs20 | -0.01970 | 0.0036 | 45 | -5.477 | <0.0001 *** |
|  |  | n_subs10 − n_subs50 | -0.01522 | 0.0036 | 45 | -4.232 | 0.0007 ** |
|  |  | n_subs10 − n_subs100 | -0.00720 | 0.0036 | 45 | -2.002 | 0.3083 |
|  |  | n_subs20 − n_subs50 | 0.00448 | 0.0036 | 45 | 1.245 | 1.0000 |
|  |  | n_subs20 − n_subs100 | 0.01250 | 0.0036 | 45 | 3.475 | 0.0069 |
|  |  | n_subs50 − n_subs100 | 0.00802 | 0.0036 | 45 | 2.230 | 0.1846 |
|  | LN | n_subs10 − n_subs20 | -0.0206 | 0.00411 | 45 | -5.024 | <0.0001 *** |
|  |  | n_subs10 − n_subs50 | -0.0654 | 0.00411 | 45 | -15.925 | <0.0001 *** |
|  |  | n_subs10 − n_subs100 | -0.0935 | 0.00411 | 45 | -22.751 | <0.0001 *** |
|  |  | n_subs20 − n_subs50 | -0.0448 | 0.00411 | 45 | -10.902 | <0.0001 *** |
|  |  | n_subs20 − n_subs100 | -0.0728 | 0.00411 | 45 | -17.727 | <0.0001 *** |
|  |  | n_subs50 − n_subs100 | -0.0280 | 0.00411 | 45 | -6.826 | <0.0001 *** |
|  | VAN | n_subs10 − n_subs20 | -0.0444 | 0.0102 | 45 | -4.357 | 0.0005 ** |
|  |  | n_subs10 − n_subs50 | -0.1042 | 0.0102 | 45 | -10.232 | <0.0001 *** |
|  |  | n_subs10 − n_subs100 | -0.1238 | 0.0102 | 45 | -12.153 | <0.0001 *** |
|  |  | n_subs20 − n_subs50 | -0.0599 | 0.0102 | 45 | -5.875 | <0.0001 *** |
|  |  | n_subs20 − n_subs100 | -0.0794 | 0.0102 | 45 | -7.796 | <0.0001 *** |
|  |  | n_subs50 − n_subs100 | -0.0196 | 0.0102 | 45 | -1.921 | 0.3665 |

| **(F) *Interaction between Scan Length and Number of Subjects (ANOVA)*** | | | | |
| --- | --- | --- | --- | --- |
| **Metric** | **Sum Sq** | **Df** | **F value** | ***p* value** |
| Edge-level ICC | 0.000677 | 9 | 0.1819 | 0.9956201 |
| Connectome-level ICC | 0.000126 | 9 | 0.6177 | 0.7649 |
| Connectome correlation | 0.00007 | 9 | 0.8459 | 0.588012 |
| Differential Identifiability | 0.000077 | 9 | 0.9836 | 0.4808 |
| Success rate | 0.6 | 9 | 0.2734 | 0.972491 |
| Discriminability | 0.00022 | 9 | 0.4810 | 0.8655 |
| Test-Retest Reliability (group maps) | 0.05178 | 9 | 1.3929 | 0.199836 |
| Test-Retest Reliability (subject maps) | 0.006118 | 9 | 0.5041 | 0.8688689 |
| Spatial Similarity (group maps) | 0.00762 | 9 | 1.2018 | 0.2928385 |
| Spatial Similarity (subject maps) | 0.001510 | 9 | 0.7546 | 0.65860 |

**Supplementary Table S3 – Effects of test–retest condition on test–retest reliability of connectivity metrics.** ANOVA results for the effect of test–retest condition (within‑ vs between‑session/run) on all node‑based and voxel‑based reliability metrics. For most metrics, the model was mean_value ~ scan_length * n_subs + session/run; for edge‑level ICC, test–retest reliability and spatial similarity, the model was mean_value ~ n_subs * scan_length * rsn + session/run. Significance levels (Bonferroni corrected for 10 comparison/metrics): *** p < 0.0001, ** p < 0.001, * p < 0.005.

| ***Main effect of Test-retest condition (ANOVA)*** | | | | |
| --- | --- | --- | --- | --- |
| **Metric** | **Sum Sq** | **Df** | **F value** | ***p* value** |
| Edge-level ICC | 0.046371 | 1 | 112.1723 | < 2.2e-16 *** |
| Connectome-level ICC | 0.000878 | 1 | 38.7056 | 1.636e-05 *** |
| Connectome correlation | 0.00010 | 1 | 10.2382 | 0.005967 |
| Differential Identifiability | 0.000001 | 1 | 0.1661 | 0.6894 |
| Success rate | 2.2 | 1 | 9.0327 | 0.008874 |
| Discriminability | 0.00577 | 1 | 113.3713 | 2.175e-08 *** |
| Test-Retest Reliability (group maps) | 0.00071 | 1 | 0.1730 | 0.678231 |
| Test-Retest Reliability (subject maps) | 0.000778 | 1 | 0.5768 | 0.4491761 |
| Spatial Similarity (group maps) | 0.01449 | 3 | 6.8589 | 0.0001701 ** |
| Spatial Similarity (subject maps) | 0.005011 | 3 | 7.5107 | 7.059e-05 *** |

**Supplementary Table S4 – Effects of resting‑state network on test–retest reliability of connectivity metrics.** ANOVA results for the effect of resting‑state network on node‑based (edge‑level ICC) and voxel‑based (test–retest reliability and spatial similarity) reliability metrics. For all metrics, the model was mean_value ~ n_subs * scan_length * rsn + session/run. Significance levels (Bonferroni corrected for 10 comparison/metrics): *** p < 0.0001, ** p < 0.001, * p < 0.005.

| ***Main effect of Resting-State Network (ANOVA)*** | | | | |
| --- | --- | --- | --- | --- |
| **Metric** | **Sum Sq** | **Df** | **F value** | ***p* value** |
| Edge-level ICC | 0.114623 | 6 | 46.2127 | < 2.2e-16 *** |
| Test-Retest Reliability (group maps) | 0.39096 | 6 | 15.7749 | 4.915e-13 *** |
| Test-Retest Reliability (subject maps) | 0.093231 | 6 | 11.5221 | 5.414e-10 *** |
| Spatial Similarity (group maps) | 0.49975 | 6 | 118.2552 | < 2.2e-16 *** |
| Spatial Similarity (subject maps) | 0.040551 | 6 | 30.3893 | < 2.2e-16 *** |

**Supplementary Table S5 – Effects of temporal downsampling strategy on test–retest reliability of connectivity metrics. (A) Main effect of temporal downsampling strategy (ANOVA).** ANOVA results for the effect of temporal downsampling strategy on all node‑based reliability metrics for a fixed sample size of 100 subjects. For most metrics, the model was mean_value ~ scan_length * strategy + session/run; for edge‑level ICC, the model was mean_value ~ scan_length * strategy * rsn + session/run. **(B) Pairwise comparisons across temporal downsampling strategies.** Post‑hoc pairwise contrasts between temporal downsampling strategies for each reliability metric corresponding to (A). **(C) Interaction between temporal downsampling strategy and scan length.** ANOVA results for the interaction between temporal downsampling strategy and scan length on reliability metrics. Significance levels (Bonferroni corrected for 10 comparison/metrics): *** p < 0.0001, ** p < 0.001, * p < 0.005.

| **(A) *Main effect of Temporal Downsampling Strategy (ANOVA)*** | | | | |
| --- | --- | --- | --- | --- |
| **Metric** | **Sum Sq** | **Df** | **F value** | ***p* value** |
| Edge-level ICC | 0.003064 | 2 | 4.0718 | 0.02056 |
| Connectome-level ICC | 0.003684 | 2 | 102.7312 | 7.639e-08 *** |
| Connectome correlation | 0.001063 | 2 | 49.1846 | 3.264e-06 *** |
| Differential Identifiability | 0.000264 | 2 | 17.973 | 0.0003419 ** |
| Success rate | 17.6 | 2 | 55.1508 | 1.847e-06 *** |
| Discriminability | 0.00470 | 2 | 75.510 | 3.759e-07 *** |

| **(B) *Pairwise comparisons across Temporal Downsampling Strategy*** | | | | | |
| --- | --- | --- | --- | --- | --- |
| ***Connectome-level ICC*** | | | | | |
| **Contrast** | **Estimate** | **SE** | **df** | **t.ratio** | **p.value** |
| Scrubbing − Segment | -0.04673 | 0.00212 | 11 | -22.069 | <0.0001 *** |
| Scrubbing − Truncation | -0.00685 | 0.00212 | 11 | -3.235 | 0.0238 |
| Segment − Truncation | 0.03988 | 0.00212 | 11 | 18.834 | <0.0001 *** |
| ***Connectome correlation*** | | | | | |
| **Contrast** | **Estimate** | **SE** | **df** | **t.ratio** | **p.value** |
| Scrubbing − Segment | -0.02699 | 0.00164 | 11 | -16.417 | <0.0001 *** |
| Scrubbing − Truncation | -0.00726 | 0.00164 | 11 | -4.418 | 0.0031 * |
| Segment − Truncation | 0.01972 | 0.00164 | 11 | 11.999 | <0.0001 *** |
| ***Differential Identifiability*** | | | | | |
| **Contrast** | **Estimate** | **SE** | **df** | **t.ratio** | **p.value** |
| Scrubbing − Segment | 8.75e-05 | 0.00135 | 11 | 0.065 | 1.0000 |
| Scrubbing − Truncation | 3.75e-04 | 0.00135 | 11 | 0.277 | 1.0000 |
| Segment − Truncation | 2.87e-04 | 0.00135 | 11 | 0.212 | 1.0000 |
| ***Success rate*** | | | | | |
| **Contrast** | **Estimate** | **SE** | **df** | **t.ratio** | **p.value** |
| Scrubbing − Segment | -1.858 | 0.199 | 11 | -9.315 | <0.0001 *** |
| Scrubbing − Truncation | -0.335 | 0.199 | 11 | -1.682 | 0.3622 |
| Segment − Truncation | 1.523 | 0.199 | 11 | 7.633 | <0.0001 *** |
| ***Discriminability*** | | | | | |
| **Contrast** | **Estimate** | **SE** | **df** | **t.ratio** | **p.value** |
| Scrubbing − Segment | -0.04312 | 0.00279 | 11 | -15.453 | <0.0001 *** |
| Scrubbing − Truncation | 0.00118 | 0.00279 | 11 | 0.421 | 1.0000 |
| Segment − Truncation | 0.04430 | 0.00279 | 11 | 15.874 | <0.0001 *** |

| **(C) *Interaction between Scan Length and Temporal Downsampling Strategy (ANOVA)*** | | | | |
| --- | --- | --- | --- | --- |
| **Metric** | **Sum Sq** | **Df** | **F value** | ***p* value** |
| Edge-level ICC | 0.005340 | 6 | 2.3659 | 0.03692 |
| Connectome-level ICC | 0.004150 | 6 | 38.5800 | 9.224e-07 *** |
| Connectome correlation | 0.002361 | 6 | 36.4057 | 1.245e-06 *** |
| Differential Identifiability | 0.000451 | 6 | 10.264 | 0.0005768 ** |
| Success rate | 9.1 | 6 | 9.5349 | 0.0007967 ** |
| Discriminability | 0.00351 | 6 | 18.761 | 3.477e-05 *** |

**Supplementary Table S6 – Effects of preprocessing pipeline on test–retest reliability of connectivity metrics.** ANOVA results for the effect of preprocessing pipeline on all node‑based and voxel‑based reliability metrics. For most metrics, the model was mean_value ~ scan_length * n_subs * pipeline + session/run; for edge‑level ICC, test–retest reliability and spatial similarity, the model was mean_value ~ scan_length * n_subs * pipeline * rsn + session/run. Significance levels (Bonferroni corrected for 10 comparison/metrics): *** p < 0.0001, ** p < 0.001, * p < 0.005.

| ***Main effect of Preprocessing Pipeline (ANOVA)*** | | | | |
| --- | --- | --- | --- | --- |
| **Metric** | **Sum Sq** | **Df** | **F value** | ***p* value** |
| Edge-level ICC | 0.01300 | 1 | 8.4095 | 0.004106 * |
| Connectome-level ICC | 0.008466 | 1 | 188.1376 | 1.044e-14 *** |
| Connectome correlation | 0.00189 | 1 | 12.4032 | 0.001352 * |
| Differential Identifiability | 0.000871 | 1 | 10.4557 | 0.0029003 * |
| Success rate | 0.6 | 1 | 1.2291 | 1.2291 |
| Discriminability | 0.01650 | 1 | 102.2278 | 2.462e-11 *** |
| Test-Retest Reliability (group maps) | 0.00068 | 1 | 0.1346 | 0.7140108 |
| Test-Retest Reliability (subject maps) | 0.000144 | 1 | 0.1309 | 0.7178671 |
| Spatial Similarity (group maps) | 0.00030 | 1 | 0.3395 | 0.5604015 |
| Spatial Similarity (subject maps) | 0.000005 | 1 | 0.0190 | 0.89051 |
